# Indoor Rewilding of Laboratory Mice Recalibrates Pulmonary Mucosal Immunity and Mechanics

**DOI:** 10.1101/2025.10.26.684616

**Authors:** Mohamed Lala Bouali, Amir Mohamed Kezai, Marie-Josée Beaulieu, Joanny Roy, Papa Yaya Badiane, Véronique Lévesque, Luc Filion, Luc Vallières, Marie-Renée Blanchet, Sébastien S. Hébert

## Abstract

Laboratory mice raised under specific-pathogen-free (SPF) conditions experience restricted microbial and antigenic exposure, which favours an immature immune system and limits their translational value for respiratory research. While microbial enrichment in “dirty” mouse models restores immune maturation, its impact on integrated respiratory function and model transferability to human disease remains understudied. Here, we tested whether ecological exposure through indoor rewilding of SPF-reared mice could reshape immune complexity and recalibrate pulmonary physiology. Two-month-old female C57BL/6J mice were housed for three months under SPF or indoor-rewilding conditions and assessed for immune, mechanical, and systemic parameters. Rewilded mice exhibited expanded pulmonary immune subsets, increased dendritic-cell immune checkpoint, with TNF/IFN-γ activation coupled to regulatory IL-10 signaling. Despite sustained exposure, the alveolar-capillary barrier integrity was preserved. Functionally, respiratory oscillometry revealed improved pulmonary mechanics, including lower airway resistance, higher compliance, and reduced airway responsiveness to methacholine. Systemic cytokine analyses indicated compartmentalized pulmonary immune activation, maintaining an overall anti-inflammatory balance. Importantly, PRIA screening detected no reportable pathogens introduced during rewilding, while cecal shotgun metagenomics confirmed microbial enrichment. Together, these findings demonstrate that indoor rewilding reestablishes coordinated lung immune and mechanical homeostasis in SPF-reared mice, providing a safe and scalable model for studying human-like mucosal immunity and respiratory physiology with broad implications for preclinical respiratory research and therapeutic testing.

## INTRODUCTION

While SPF mice provide the controlled, infection-free models essential for reproducible research, this very cleanliness imposes a significant biological constraint: a poorly trained immune system. In contrast to humans, who encounter a diverse array of microbes and antigens, SPF mice develop limited lymphocytic and inflammatory profiles, leading to distorted infection responses and compromised translational fidelity of findings from murine disease models [1,2]. To bridge this gap, so-called “dirty” mouse models have emerged, which deliberately reintroduce ecological complexity. Approaches such as embryo transfer into wild dams (“wildlings”), co-housing with feral mice, and rewilding in secure outdoor enclosures have been shown to restore many hallmarks of immune maturity, including expanded granulopoiesis, more robust antibody and cytokine responses, and human-like patterns of pathogen susceptibility [3–7]. However, these “naturalization” paradigms remain logistically complex, seasonally constrained, and carry risks of zoonotic transmission, alongside varying effects depending on the exposure setup, thus restricting their scalability and reproducibility [8,9].

The lungs, being in constant contact with the external environment, are particularly affected by the SPF conditions. Respiratory resilience, i.e., the ability of the lung tissue to resist, adapt, and recover from insults while maintaining efficient gas exchange, requires coordinated epithelial-endothelial integrity, immune surveillance, and mechanical performance, all of which are shaped by environmental exposure. SPF housing, by shielding animals from microbial, antigenic, and particulate challenges (e.g., allergens), profoundly limits the maturation of pulmonary immunity. The scarce presence of lung parenchymal mast cells in SPF mice [10], in clear contrast to their abundant presence in human lungs [11,12], exemplifies the simplified leukocyte repertoires in laboratory mice that would translate into unrealistic mimicking of human lung physiology and pathology [13]. Interestingly, recent studies in “dirty” mouse models have showed that ecological exposure induces pulmonary immune remodeling, including recruitment of parenchymal mast cells akin to feral mice [14], and modulation of allergen-induced hyperresponsiveness, which can be either heightened [15] or attenuated [16] depending on the exposure setup. However, whether such ecological enrichment also remodels pulmonary properties, particularly its mechanical performance, has never been tested.

In this study, we developed a novel indoor rewilding paradigm (i.e., an indoor semi-natural, habitat-based vivarium) within a non-SPF animal facility that combines high ecological complexity, controllable experimental parameters, and year-round operational convenience. This semi-natural vivarium replicates key elements of feral mouse habitat (e.g., farmland soil, vegetation, invertebrates, manure, and organic substrates), whose composition and renewal are precisely controlled, while maintaining standardized conditions for temperature, humidity, light cycles, and predator exclusion. This paradigm allows to study how ecological exposure recalibrates host physiology in a reproducible, biosecure, and scalable setting, while promoting animal welfare through natural behaviors like foraging and burrowing.

Leveraging this platform, we conducted the first comprehensive assessment of pulmonary and systemic immunity, barrier integrity, and partitioned respiratory mechanics in SPF-reared mice after three months of indoor rewilding. It is well established that lung function is closely coupled to immune activity, and that persistent immune activation and parenchymal alterations drive the progressive decline in pulmonary performance characteristic of chronic respiratory diseases such as asthma and chronic obstructive pulmonary disease (COPD) [17]. Hence, we sought to determine whether ecological immune enrichment could translate into measurable changes in lung mechanical function. Our results demonstrate that indoor rewilding diversifies the pulmonary immune landscape while enhancing mechanical efficiency and maintaining barrier integrity, a combination not previously characterized in “dirty” mouse models. These findings position indoor rewilding as a complementary and translational tool for modelling human-like mucosal immunity and pulmonary physiology, with broad implications for respiratory disease research and preclinical therapy development.

## MATERIALS AND METHODS

### Ethics statement

All procedures involving mice were carried out in accordance with the Canadian Council on Animal Care (CCAC) guidelines and were approved by the Animal Care Committee (CPAUL-2, approval number: 23-1342). Female mice were chosen to reduce confounding factors related to breeding and limit aggression within the vivarium.

### Animals

Two-month-old female C57BL/6J mice were purchased from Jackson Laboratories (strain #000664). Upon arrival, mice were randomly assigned to standard SPF housing or to an indoor semi-natural vivarium. Both groups were maintained at ∼22°C, ∼45%RH, on a 12:12-hour light/dark cycle, and had *ad libitum* access to food and water.

### Semi-natural vivarium

Our indoor rewilding platform consisted of a 12m^2^ enclosure (adapted swimming pool) (see **Fig. 1A; Supp. Fig. 1; Supp. Video 1**) established within a non-SPF room ordinarily dedicated to farm animals at the IUCPQ Research Center animal facility (Québec-City, Canada). Before initiating the experimental protocol, 28 two-month-old female mice were randomly allocated to SPF (14) or Rewild (14). Afterwards, all mice were ear-tagged with ear punches, and baseline health status and body weight were recorded. Both groups were comparable at baseline, with no differences in health or weight. No animals were excluded from this study. SPF group remained in conventional SPF housing in individually ventilated cages (IVCs). The Rewild group was released into the semi-natural vivarium for a three-month rewilding period, with continuous infrared video monitoring (**Supp. Video 2**). Rewilded mice received standard laboratory chow and scattered bird seeds. After the rewilding period, mice were captured using Longworth traps, then transferred to non-ventilated standard cages (not IVCs) containing vivarium derived materials (e.g., soil, plants) and were maintained on same diet as in the vivarium, to maintain the same ecological exposure until testing. Rewild and SPF groups were weighed and examined prior to experimentation (no mortality or abnormal phenotypes were detected). To minimize stress post-capture, experimental testing began 24h after the last mouse of each cohort was captured, resulting in a 24-48h holding period.

**Figure 1.**
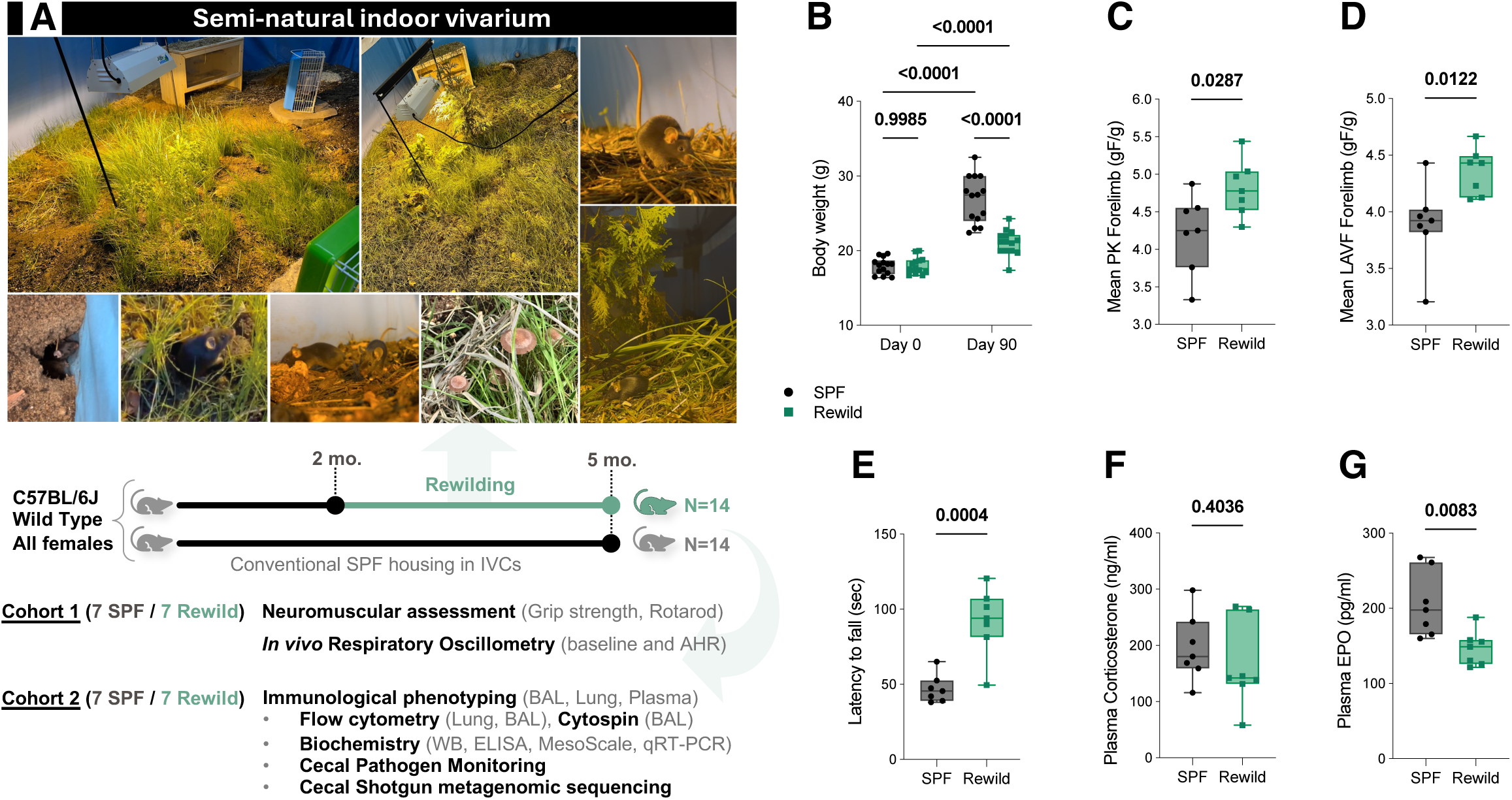
Rewilding promotes the maintenance of a lean body weight and enhances neuromuscular and systemic performance. **(A)** Representative images of the semi-natural indoor vivarium and schematic of the experimental design (see **Supp. Fig. 1**). Two-months-old female C57BL/6J mice were housed under SPF or rewilding (semi-natural vivarium) conditions for 90 days. Complete recapture required 4 trapping nights, with a 100% recapture rate (14/14). Traps were checked regularly via cameras limiting time in-trap to ≤11 h; Longworth traps contained nesting material, food (peanut butter, seeds, chow), and water gel. Cohort 1 (N = 7/group) was used for neuromuscular testing (grip strength, rotarod) and pulmonary mechanics using forced oscillation technique (FOT); while cohort 2 (N = 7/group) for immunophenotyping, biochemical assays, and cecal pathogen monitoring and shotgun metagenomics. All animals were analyzed; no exclusions or outliers were removed from either group. **(B)** Body weight at baseline (day 0) and endpoint (day 90). **(C-D)** Forelimb grip strength was assessed as peak force (PK) and last available force (LAVF), normalized to body weight. **(E)** Neuromotor coordination and endurance assessed by latency to fall on the rotarod. **(F)** Plasma corticosterone and **(G)** erythropoietin (EPO) levels measured by ELISA. Data are shown as individual values overlaid on box-and-whisker plots, where the box spans the interquartile range (25^th^-75^th^ percentile), the line indicates the median, and whiskers extend from the minimum to the maximum. Statistical comparisons were performed using two-way repeated measures ANOVA with Šidák’s multiple comparison test for body weight, and unpaired two-tailed Student’s t-tests for all other comparisons.

### Outcome measures and sample size determination

This study aimed to characterize respiratory, immune, and molecular adaptations to indoor rewilding. No single primary outcome was predefined to optimize the utilization of animals and tissue samples. Sample size was set at N=7 per group, based on pilot studies and our extensive experience with the FlexiVent system and flow cytometry, where this number consistently detected meaningful differences in respiratory mechanics and immune phenotype, respectively. Using females only reduced sex-related variability in lung mechanics and immunity, further supporting statistical power at this N. The high sensitivity and low variability of FlexiVent allowed robust detection with minimal animal use, in line with the 3Rs principle. The same group size was used for immunological assays to ensure consistency and cross-validation. Age, sex, temperature, humidity, light cycle, circadian phase and handling (with alternating group testing), and experimenter consistency (same experimenter throughout) were standardized to minimize confounders and ensure experimental reproducibility. No blinding was performed during experimentation or data analysis.

### Muscular force and motor coordination testing

At endpoint (5-month-old mice), seven mice per group (randomly selected) (cohort 1, also used for FlexiVent, see **Fig. 1A**) underwent grip strength and rotarod testing during the light phase in a dedicated room after a 30-min acclimation. Forelimb grip strength was measured in a single session following the modified SHIRPA and using a digital force meter (Ametek® Chatillon Columbus Instruments), as indicated before [18]. Each mouse performed three trials (1-min intervals); peak force (PK) and last available force (LAVF) were recorded and normalized to body weight. Neuromotor coordination and endurance were evaluated using a Rotarod apparatus (Panlab, Harvard Bioscience). Mice were trained on two consecutive days (4 RPM for 5-min). On the third day, mice underwent two testing trials (30-min inter-trial interval) with linear acceleration from 4 to 40 RPM over 300-sec; the latency-to-fall was averaged.

### *In vivo* respiratory mechanics assessment

Partitioned oscillometry was used to assess *in vivo* respiratory mechanics in mice *via* the FlexiVent^®^ system (FX module 2, FlexiWare version 8.3, SCIREQ, Montréal, Canada), following well-established protocols [19–23].

### Mechanical ventilation

Mice were anesthetized by intraperitoneal injection of ketamine (100mg/kg) and xylazine (10mg/kg). At surgical depth of anesthesia, a tracheostomy was performed, and an 18-gauge metal cannula secured with suture. Mice were ventilated supine at 10mL/kg tidal volume, 150 breaths/min, PEEP 3cmH₂O, with room air as carrier gas. Pancuronium bromide (100μL at 20μg/ml, intramuscular) suppressed spontaneous respiratory efforts. Anesthetic depth was monitored throughout the experiment using electrocardiography.

### Baseline lung mechanics

Two deep inflations to 30cmH_2_O were performed before data collection [24]. Baseline respiratory mechanics were then assessed through three cycles (1-min intervals) of perturbations sequence: a Deep-inflation (yielding inspiratory capacity, IC), followed by Snapshot-150, Quick Prime-3, and pressure-controlled partial pressure-volume (PVs-P) maneuvers. The Snapshot-150 maneuver (1.25-sec sinusoidal oscillation at 2.5Hz, single-frequency forced oscillation technique, FOT) provided single-compartment estimates of respiratory system resistance (Rrs), elastance (Ers), and the normalized work of breathing (WOBn). The Quick Prime-3 multifrequency perturbation (1-20Hz, broadband FOT) was used to fit the constant-phase model and derive Newtonian resistance (Rn), tissue damping (G), tissue elastance (H), and hysteresivity (η=G/H) [25]. PVs-P loops were recorded using a stepwise inflation to 30cmH_2_O, yielding the Salazar-Knowles shape parameter K (volume-independent compliance). Measurements with coefficient of determination (COD) <0.9 were excluded; remaining values were averaged per mouse.

### Methacholine challenge

After baseline recordings, increasing concentrations of aerosolized methacholine (0, 3, 10, 100, 300mg/ml in PBS; Sigma-Aldrich) were delivered for 10-sec at 50% duty cycle via an integrated Aeroneb Lab nebulizer (rinsed with PBS between doses) [19,23]. To minimize derecruitment, two deep-inflations to 30cmH_2_O preceded each dose. After each nebulization, alternating Snapshot-150 and Quick Prime-3 perturbations were repeated 8-10 times, with ∼8 sec of tidal ventilation between maneuvers. Measurements with COD<0.9 were excluded. For each dose, the maximal post-dose values (Δ from dose 0mg/ml baseline) for Rrs, Ers, Rn, G, H, and WOBn were used to draw the concentration-response curves, and compute AUCs.

### Bronchoalveolar lavage (BAL)

In cohort 2, mice under deep ketamine/xylazine anesthesia were tracheotomized, and a bronchoalveolar lavage was performed by doing three sequential 1mL PBS injections/recoveries [26,27]. Recovered volumes were pooled and recorded. Samples were centrifuged (500g, 5 min, 4°C). Cell pellets were resuspended in 200μL PBS for total/differential counts and flow cytometry, while BAL fluid (BALF) were aliquoted and stored at -80°C for albumin ELISA and MSD multiplex cytokines/chemokines. Total BAL cell counts used an automated cell counter (Corning CytoSmart Cell Counter, Axion AxISVue, Cell-Count v3). Cytospins (Cytospin III, Shandon Scientific) were stained (Hema-3, Thermo Fisher, #122-911) for differential counts under light microscopy.

### Blood collection and plasma preparation

Blood was obtained by cardiac puncture into EDTA tubes and centrifuged (2000g, 10 min, 4°C). Plasma was aliquoted and stored at -80°C until use for ELISAs (erythropoietin, corticosterone) and MSD multiplex cytokines/chemokines.

### Lung tissue processing and leukocyte isolation

Lungs were excised, and left, right superior, and post-caval lobes were processed fresh and finely minced in ice-cold GNK buffer (PBS supplemented with glucose, NaCl, and KCl), then digested for 45 min at 37°C with gentle inversion every 10 min in equal volume of GNK buffer containing 1mg/ml collagenase IV (Worthington, #LS004188), 0.2mg/ml DNase I (Sigma, #10104159001), and 10% FBS (Gibco). Samples were then mechanically dissociated through a 70-µm cell strainer. Following PBS wash and centrifugation (500g, 5 min, 4°C), RBC lysis (ammonium chloride solution) was performed for 4 min at room temperature, quenched with PBS, and suspension filtered (40-µm). Final pellets were resuspended in PBS containing 1% FBS and counted using an automated cell counter, excluding dead cells stained with trypan blue. For fluorescence minus one (FMO) controls, pooled samples were used. This well-established protocol yields robust recovery of viable CD45^+^ cells for downstream multiparametric flow cytometry [26–29].

### Flow cytometry

BAL cells and lung single-cell suspensions were stained with fluorochrome-conjugated antibodies (**Supp. Table 1**). Data were acquired on a BD LSR Fortessa (BD Biosciences) and analyzed using FlowJo v10. Gating strategies were based on FMO controls and are provided in **Supp. Figs. 2-5**.

### RNA extraction and quantitative RT-PCR

Total RNA from frozen right-middle lobes was extracted (TRIzol reagent), quantified (Infinite-2000, TECAN), reverse-transcribed (High-Capacity cDNA kit, Thermo-Fisher, #4368814), and amplified using SYBR chemistry (Thermo-Fisher, #A46109). Primers (**Supp. Table 2**) were designed with melting temperatures around 60°C, and their specificity verified *in silico* (Primer-BLAST, NCBI). Amplifications were performed on a LightCycler 480 II system (Roche) using the following cycling conditions: 50°C for 2 min, 95°C for 10 min, followed by 40 cycles of 95°C for 30 sec and 60°C for 30 sec, and a final melting curve analysis (95°C for 1 min, 55°C for 30 sec, 95°C for 30 sec). Expression was normalized to RPLP0 and calculated by 2^(−ΔΔCt).

### Protein extraction and immunoblotting

Frozen right-inferior lobes were homogenized in RIPA lysis buffer containing protease and phosphatase inhibitors (Roche). Equal amounts of protein (30µg) were resolved by SDS-PAGE and transferred onto PVDF or nitrocellulose. Membranes were blocked with 5% non-fat dry milk in TBS-Tween and probed overnight at 4°C with primary antibodies, followed by HRP-conjugated secondary antibodies, and detected by ECL (Immobilon® ECL UltraPlus Western HRP Substrate). Details regarding all antibodies are provided in **Supp. Table 3**. Total proteins were used for normalization (No-Stain reagent on PVDF, Ponceau-S on nitrocellulose). Densitometry used ImageJ, and representative lanes from the immunoblots are displayed for each protein of interest.

### Enzyme-linked immunosorbent assays (ELISA)

Commercial ELISA kits were used according to manufacturer instructions for quantification of plasma erythropoietin (EPO, Abcam, #Ab270893), plasma corticosterone (Biotechne, #NBP3-23556), and BALF albumin (Abcam, #207620). Results were normalized to appropriate standard curves.

### Multiplex MSD cytokines/chemokines profiling

BALF and plasma cytokines/chemokines were quantified using a V-PLEX Mouse Cytokine 19-Plex Kit (MSD) following the manufacturer’s protocol. Plates were read on a MESO^®^ QuickPlex SQ-120 Multiplex Imager. Data were analyzed using MSD-Workbench software. Results were log_10_-transformed prior to statistical analysis to account for skewed distributions.

### Cecal pathogen monitoring

Under a Class II biosafety cabinet, cecal contents were collected aseptically from SPF and rewilded mice using sterile instruments and DNase/RNase-free tubes. Samples were snap-frozen and stored at -80°C until shipment on dry ice to Charles River Research Animal Diagnostic Services (RADS). Pathogen screening was performed using the PCR Rodent Infectious Agent (PRIA^®^) panels. Results are summarized in **Supp. Table 4**.

### Cecal Shotgun metagenomic sequencing

DNA extraction, library preparation, sequencing, and primary processing of cecal contents were performed by Microbiome Insights (Vancouver, Canada). Paired-end sequencing (2x150bp) was run on an Illumina NextSeq 2000. Reads were processed using the Sunbeam pipeline, including high-quality filtering and host/contaminant removal by mapping to the mouse genome (Mus musculus, GRCm38.p6) and the human genome. Remaining reads were taxonomically classified using Kraken2 (PlusPF database), and functional profiles were generated against SEED Subsystems (levels 1-3). Shotgun metagenomics was performed on a subset of mice (n=6/group) randomly selected from the cohort 2 (n=7/group).

### Statistical analyses

GraphPad Prism v10 was used for statistical analyses. Data distribution was assessed with the Shapiro-Wilk normality test. For comparisons between two independent groups, unpaired two-tailed Student’s t-tests were used for normally distributed data, while Mann-Whitney tests were applied for non-parametric data. For body weight trajectories, methacholine challenge curves and cytokine log-transformed values, repeated-measures two-way ANOVA was performed with Šidák’s *post hoc* correction for multiple comparisons. AUCs across doses were compared by unpaired two-tailed t-tests. Shotgun taxonomy and functions data were analyzed in R software v4.5.2 using a compositional data analysis (CoDA) framework. Relative abundances in feature tables (phylum/genus/species/functions) were TSS-normalized, zeros were replaced using the multiplicative simple replacement strategy (zCompositions package) followed by a Centered Log-Ratio (CLR) transformation (compositions package). Beta-diversity was assessed using Aitchison distances and tested by Permutational Multivariate Analysis of Variance (PERMANOVA, 924 permutations) using the vegan package. Differential abundance (DA) was assessed on CLR-transformed features (filtered as indicated) using two-sided Welch’s t-tests, and Benjamini-Hochberg FDR correction. Significance threshold for all analyses was set at p<0.05.

## Supporting information

Revised Supplementary Materials

## Patient and public involvement

Patients and/or the public were not involved in the design, conduct, reporting, or dissemination plans of this research.

## Funding statement

This work was supported by the Canadian Institute of Health Research (CIHR), the Natural Sciences and Engineering Research Council of Canada (NSERC), and the *Fonds de Recherche du Québec en Santé* (FRQS).

## Data availability

The datasets generated in the current study are available upon request.

## Acknowledgements

We thank all family members of *La ferme du vieux caveau* farm located in Saint-Joachim, Québec, for generously providing soil and other organic materials for our indoor enclosures. We also thank Justin Robillard and all staff members of the IUCPQ animal facility for their support and guidance. Our gratitude extends further to the CRCHUQ animal facility team, particularly to Marie-Claude Richer, for their continued assistance. Finally, we are grateful to Dr. Andrea L. Graham from Princeton University, New Jersey, USA, for her valuable insights and recommendations on implementing an indoor rewilding system.

## RESULTS

### Rewilding preserves a lean body weight trajectory and improves neuromuscular performance

We first compared the physiological status of two-month-old female C57BL/6 mice obtained from The Jackson Laboratory, housed for 90 days either in an SPF facility or in the semi-natural vivarium (**Fig. 1A**). During this period, no mice contracted pathogens (**Supp. Table 4**), as revealed by PCR testing of feces designed to screen for a range of viruses, bacteria, and parasites, supporting that our enclosures and procedures were biosecure.

Body weight gain was significantly different between the two groups (**Fig. 1B**). At baseline (day 0), body weights were similar (SPF: 18.05 ± 0.70 g; Rewild: 17.93 ± 0.65 g; p=0.98). After three months, SPF mice had gained 3 times more weight (27.11 ± 0.80 g; Δ +9.1, p<0.0001) than rewilded mice (21.13 ± 0.63 g; Δ +3.2, p<0.0001). A mixed-effects analysis confirmed a significant time-group interaction (F(1,30) = 62.8, p<0.0001), indicating that body weight gain was strongly influenced by housing conditions. At necropsy, abdominal fat was frequently observed in SPF mice, but not in rewilded mice (not shown).

Neuromuscular performance consistently improved under rewilding conditions. Rewilded mice appeared more vigorous and active in the vivarium (**Supp. Videos 1-2**). Forelimb grip strength was significantly higher for both peak force (**Fig. 1C**) and last available force (**Fig. 1D**). Motor coordination and endurance, assessed by rotarod latency-to-fall, were also enhanced in rewilded mice (**Fig. 1E**), indicating better coordination and vigilance. Overall, rewilded mice resisted excess weight gain and performed better functionally, contrasting with the overweight and reduced activity of SPF mice.

Plasma endocrine profiling offered complementary systemic readouts. Stress levels were evaluated by measuring corticosterone at the morning circadian trough, which did not differ between groups (**Fig. 1F**), indicating no chronic activation of the stress axis and no persistent stress response detectable after handling at the end of the rewilding period. Conversely, circulating erythropoietin (EPO) was significantly lower in rewilded mice (**Fig. 1G**). Since EPO secretion reflects renal hypoxia sensing, this decrease is consistent with lower basal hypoxic signaling. However, given that EPO can be modulated by factors such as physical activity, hematocrit, or hormonal influences, these data should be interpreted as descriptive rather than mechanistic.

Taken together, these results demonstrate that indoor rewilding is safe, non-stressful, and beneficial for the physical and metabolic fitness.

### Rewilding remodels baseline lung mechanics and blunts airways hyperresponsiveness

We next examined the effects of rewilding on basal lung mechanics and airway responsiveness to methacholine challenge using partitioned oscillometry and quasi-static pressure-volume (PVs-P) analyses with the FlexiVent system (**Fig. 2A**).

**Figure 2.**
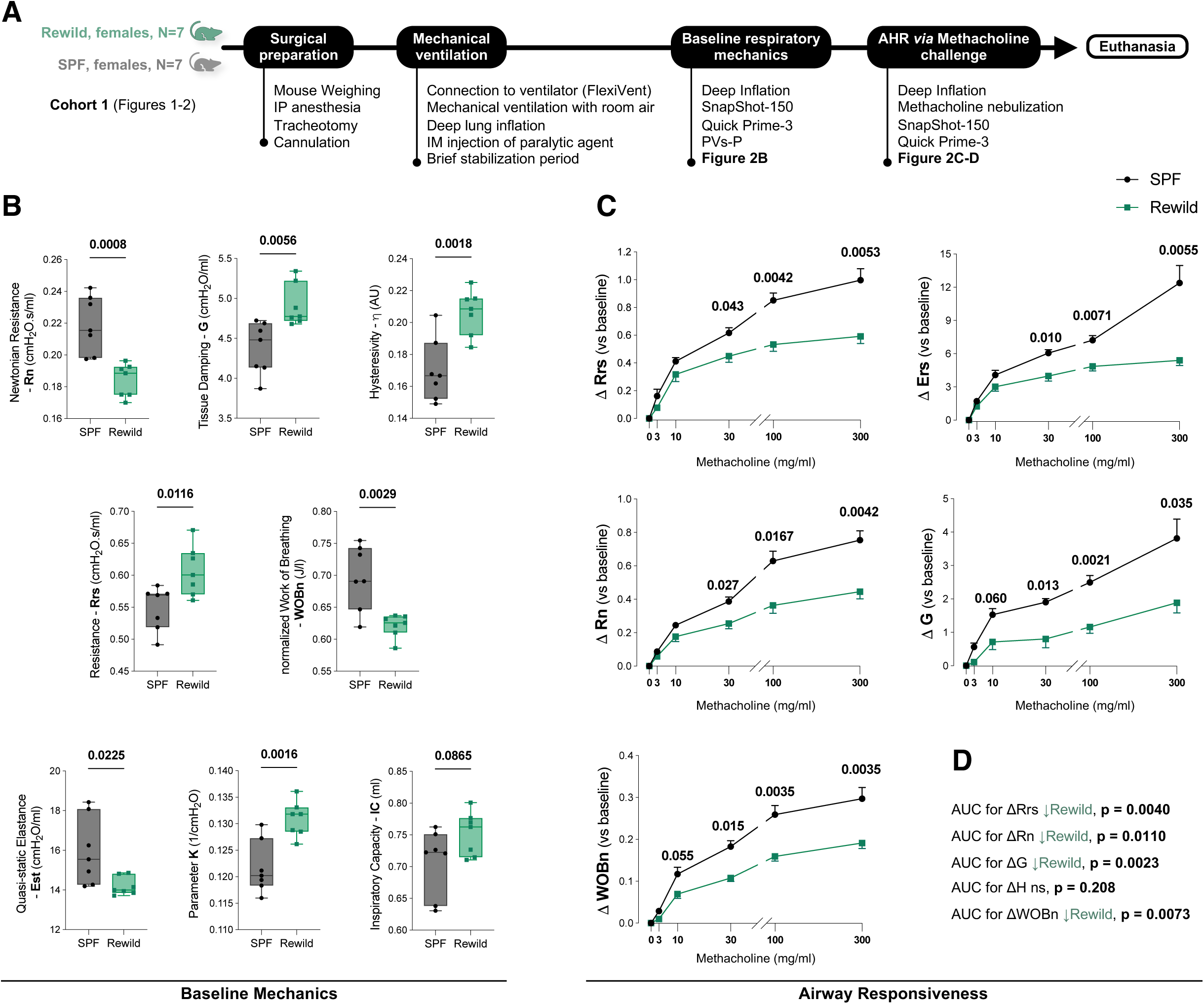
Rewilding remodels baseline lung mechanics and blunts airway responsiveness. **(A)** Experimental workflow for *in vivo* pulmonary function testing by oscillometry (FOT) using the FlexiVent system, including baseline assessment of lung mechanics and methacholine challenge to evaluate airway hyperresponsiveness (AHR). **(B)** Baseline FlexiVent readouts obtained from Deep-inflation, Snapshot-150, Quick Prime-3 and pressure-controlled partial pressure-volume (PVs-P) maneuvers. Each value represents the mean of three valid cycles per animal; comparisons between SPF and rewilded groups were made using unpaired two-tailed Student’s t-tests. **(C)** FlexiVent dose-response curves to nebulized methacholine (0-300 mg/ml) showing changes (Δ vs. baseline) in Rrs, Ers, Rn, G, and WOBn. Curves show group means of peak responses (N = 7/group); statistical comparisons were performed by two-way repeated measures ANOVA with Šidák’s correction for multiple comparisons. **(D)** Area under the curve (AUC) analyses confirm blunted AHR in rewilded mice for ΔRrs, ΔRn, ΔG, and ΔWOBn, with no significant difference for ΔH. Means were compared using unpaired two-tailed Student’s t-test. No animals or data points were excluded from analysis.

At rest, partitioned oscillometry revealed a distinct mechanical profile in rewilded mice compared to SPF controls (**Fig. 2B**). Newtonian resistance (Rn), which indexes the resistance of the central and large conducting airways, was significantly lower in rewilded mice, consistent with reduced airway tone or larger conducting calibers. In contrast, tissue damping (G), a measure of energy dissipation within the lung parenchyma, was higher, while tissue elastance (H), a proxy for parenchymal stiffness, remained unchanged (p=0.1881, not shown). Consequently, hysteresivity (η=G/H), which quantifies the fraction of elastic energy lost during each breathing cycle and serves as indicator of ventilation heterogeneity, was increased. The total respiratory resistance (Rrs), which conflates both airway and tissue components, was modestly elevated, because the rise in G outweighed the fall in Rn. Importantly, the normalized work of breathing (WOBn), which indexes the energy required to inflate the lungs, was reduced in rewilded mice, indicating a lower energetic cost of ventilation despite increased parenchymal dissipation.

Quasi-static PVs-P analysis further supported enhanced distensibility. Static elastance (Est), representing the overall stiffness of the respiratory system, was significantly lower (**Fig. 2B**), with a concomitant rise in the parameter K, a volume-independent proxy of compliance. Together, these changes indicate more compliant lungs with improved alveolar recruitment at low pressures in rewilded mice, independent of body size or weight. Inspiratory capacity (IC) showed a trend toward elevation (p=0.0865). Overall, this combination of freer central airways, higher parenchymal compliance, and slightly increased damping supports more efficient ventilation and indicates that rewilded lungs operate in a state of improved mechanical efficiency and airway recruitment. This mechanical heterogeneity may reflect localized immune-tissue interactions, as supported by the immunological findings described later.

Given that airway reactivity is altered in lung disease such as asthma and is often tested in SPF-based mouse models, we tested whether the baseline phenotype observed in **Fig. 2B** altered airway responsiveness to constrictive stress. Following increased, non-cumulative, doses of aerosolized methacholine (0 to 300mg/ml), SPF mice exhibited a steep, dose-dependent increase in airway and tissue resistive parameters, whereas rewilded mice showed significantly blunted responses across intermediate and high doses, revealing a protective phenotype with blunted airway responsiveness across both central and peripheral compartments (**Fig. 2C**). The rise in ΔRrs and ΔErs was markedly attenuated in rewilded mice across multiple doses (adjusted tests, p≤0.043 at 30 to 300mg/ml). Partitioning this response revealed reduced large-airway reactivity (ΔRn, p≤0.027 at 30 to 300mg/ml) and diminished tissue damping (ΔG, p≤0.013 at 30 to 300mg/ml). Consistently, the rise in ΔWOBn was also dampened in rewilded mice (p≤0.015 at 30 to 300mg/ml). By contrast, ΔH increased with dose in both groups without any difference (not shown), indicating that rewilding selectively protected against resistive rather than elastic loading. Consistent with decreased sensitivity to methacholine, rewilded mice exhibited significantly lower AUCs for ΔRrs, ΔRn, ΔG and ΔWOBn, but not for ΔH (**Fig. 2D**).

Together, these data demonstrate that rewilded mice exhibit reduced central airway resistance, higher compliance, and attenuated airway responsiveness, indicating a mechanically more efficient and less reactive lung compared to SPF controls.

### Rewilding densifies and reshapes lung immune compartment

We next examined whether functional remodeling of the lung was accompanied by changes in immune composition (**Fig. 3A**). Cytospin differentials showed selective increases in lymphocytes, macrophages, and neutrophils, while eosinophils remained unchanged (**Fig. 3B-C**). This profile indicates an enriched but non-eosinophilic milieu, consistent with a diversified leukocyte landscape without allergic-type profile. Flow cytometry of BAL cells confirmed these findings (**Fig. 3D**). Counts of CD45^+^ cells per BAL volume were higher in rewilded mice, with expansions of CD4^+^ and CD8^+^ T cells, B cells, and natural killer T cells (NKT).

**Figure 3.**
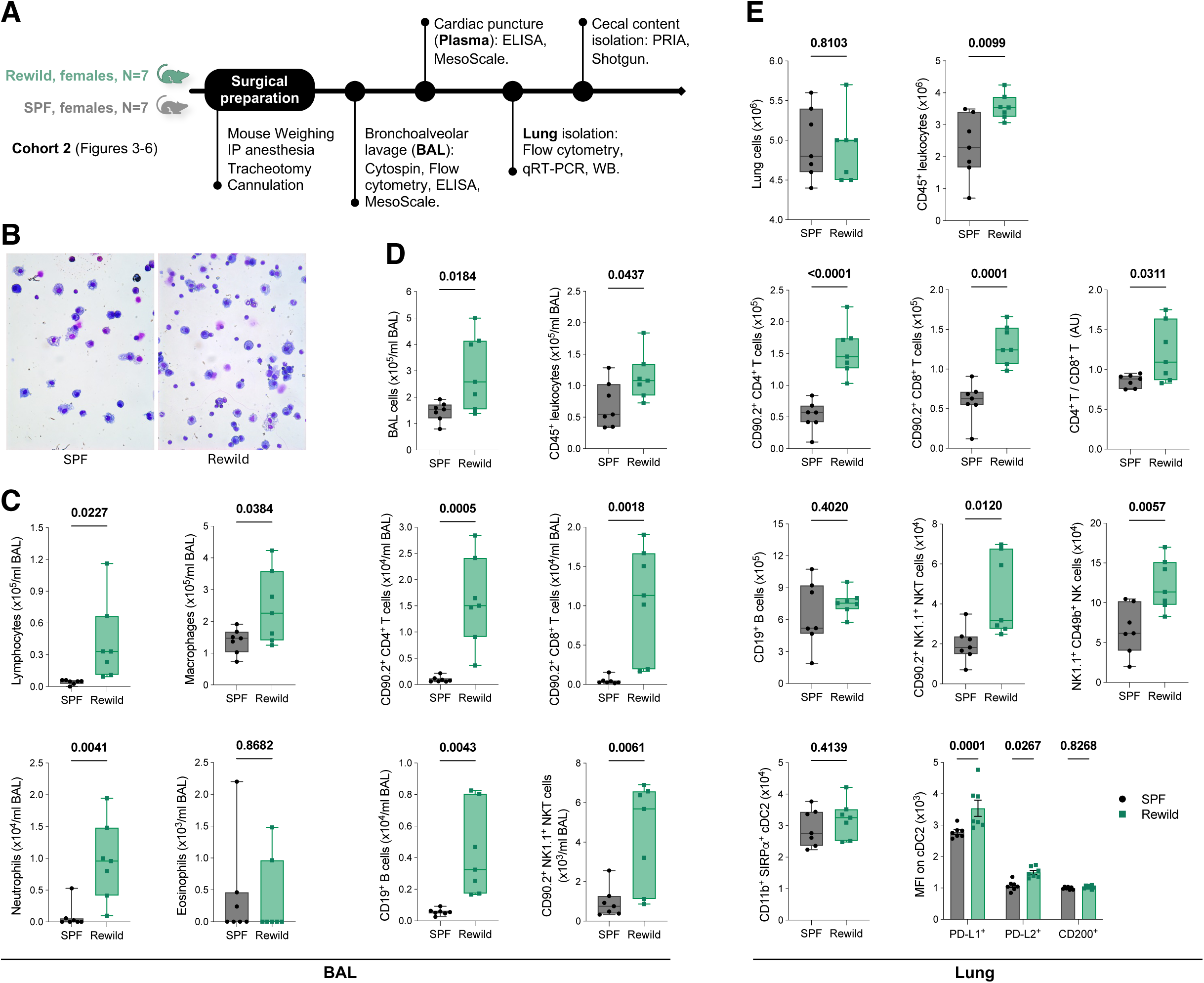
Rewilding densifies airway leukocytes and reshapes lung immune compartments. **(A)** Experimental workflow for immunophenotyping of bronchoalveolar lavage (BAL), lung and plasma compartments. **(B)** Representative cytospin images of BAL cells stained with modified May-Grünwald–Giemsa. **(C)** Differential BAL cytology showing increased lymphocytes, macrophages, and neutrophils but unchanged eosinophils in rewilded mice. **(D)** BAL flow cytometry showing higher leukocyte counts and expansions of CD4⁺ and CD8⁺ T cells, B cells, and NKT cells. **(E)** Lung single-cell suspensions showing higher leukocyte numbers and selective rewilding-increases in CD4⁺, CD8⁺, NK, and NKT cells, with stable total cell counts and B cells. Median fluorescence intensity (MFI) analysis of lung conventional cDC2 subset revealed upregulation of PD-L1 and PD-L2, but not CD200. Data are shown as individual values or as mean ± SEM for MFI histograms. Means were compared using unpaired two-tailed Student’s t-test. No animals or data points were excluded from analysis.

CD45^+^ leukocytes were significantly higher in rewilded lungs (**Fig. 3E**). Within this expanded leukocyte pool, CD4^+^ and CD8^+^ T cells were significantly increased, with a modestly higher CD4/CD8 ratio (p=0.0311) suggesting a shift toward a regulatory-dominant immune state compatible with enhanced immune surveillance and mucosal tolerance in rewilded lungs. Further, natural killer cells (NK) and NKT cells were also significantly increased, while B cells remained unchanged (**Fig. 3E**). Notably, type-2 conventional dendritic cells (cDC2s; identified as CD11c^+^ MHCII^+^ CD11b^+^ SIRPα+; see **Supp. Fig. 5 and Supp. table 1**) were similar between groups, but their phenotype shifted toward higher PD-L1 (MFI, p=0.0001) and PD-L2 (MFI, p=0.0267) expression in rewilded mice, whereas CD200 expression was unchanged (**Fig. 3E**). These results demonstrate selective enrichment of lymphoid and innate-like populations in the airspaces and lung parenchyma, accompanied by increased checkpoint tone on dendritic cells, suggestive of heightened immune surveillance without eosinophilia.

### Rewilding calibrates lung inflammatory-regulatory programs and enhance systemic anti-inflammatory balance

At the transcriptional level, rewilded lungs showed higher expression of TNF, CCL4, CXCL10, and IL-10 compared to SPF controls (all p≤0.03), while IL-4, IL-5, IL-6, INF-γ, CXCL15, and TGF-β1 remained unchanged (**Fig. 4A**). This pattern suggests a mixed but balanced state coupling effector readiness (TNF, CCL4, CXCL10) with local regulation (IL-10). At the protein level, rewilded lungs showed increased platelet endothelial cell adhesion molecule-1 (PECAM-1) and intercellular adhesion molecule-1 (ICAM-1), while vascular cells adhesion molecule-1 (VCAM-1) remained unchanged, compatible with enhanced but controlled leukocyte trafficking in the lung microvasculature (**Fig. 4B**).

**Figure 4.**
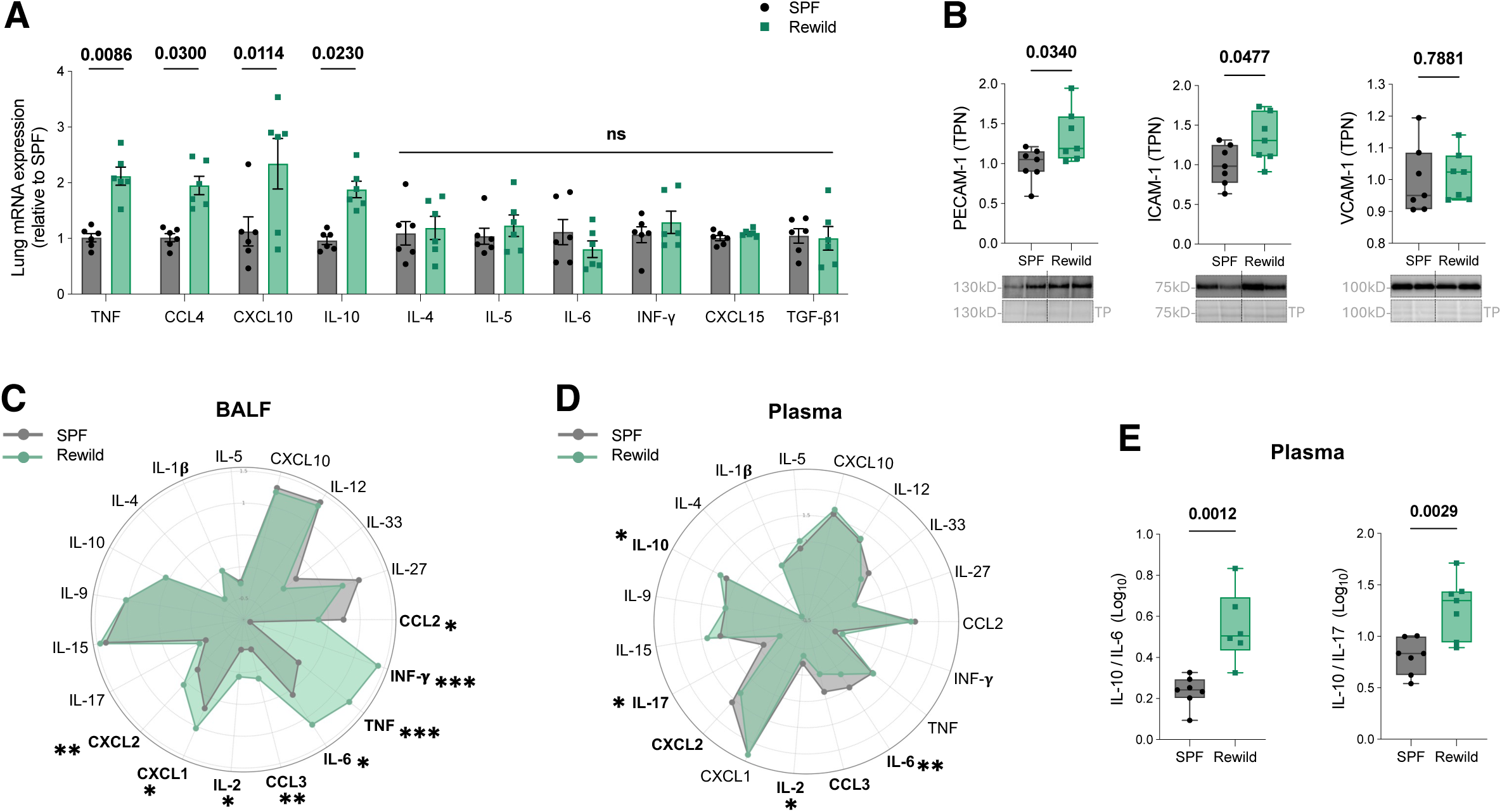
Rewilding calibrates lung inflammatory-regulatory programs with localized airway activation and systemic restraint. **(A)** Relative mRNA expression of inflammatory and regulatory genes in lung tissue measured by qRT-PCR, normalized to the housekeeping gene RPLP0. Statistical analysis was performed on Log_2_-transformed values using two-way ANOVA with Šidák’s correction for multiple comparisons. Adjusted p-values are shown; ns: non-significant. **(B)** Immunoblot analysis of vascular adhesion proteins PECAM-1, ICAM-1 and VCAM-1 in whole-lung homogenates. Representative blots and total protein normalization controls are shown; Comparisons were made using unpaired two-tailed Student’s t-tests. **(C)** Radar plot of bronchoalveolar lavage fluid (BALF) cytokines and chemokines measured by multiplex MesoScale assay. Values represent group means of log_10_-transformed concentrations (N = 7/group). The rewild group displayed broader cytokine activation in the airways, consistent with localized immune enrichment. Asterisks (*) indicate statistically significant differences between groups (unpaired two-tailed Student’s t-tests. *p < 0.05, **p < 0.001, ***p < 0.0001). **(D)** Radar plot of plasma cytokines and chemokines measured by multiplex MesoScale assay (similar analysis as in BALF, N = 7/group) illustrating a restrained profile in rewilded mice. Panels **(C)** and **(D)** are shown side by side to enable direct comparison of local (airway) versus systemic cytokine landscapes. **(E)** Ratios of systemic cytokines (log_10_-transformed), IL-10/IL-6 and IL-10/IL-17, highlighting increased systemic anti-inflammatory balance. Means were compared using unpaired two-tailed Student’s t-test. No animals or data points were excluded from analysis.

Further analysis of circulating cytokines/chemokines in BALF revealed a selective pro-inflammatory enrichment under rewilding (**Fig. 4C**). The radar plot highlighted elevated IFN-γ, TNF, CXCL1, CXCL2, and CCL3, IL-2 and IL-6 (all p≤0.0454, individual t-tests), consistent with an activated but non-allergic pulmonary milieu. Furthermore, plasma levels of the same mediators suggest that systemic inflammation is not present in rewilded mice. Plasma cytokine radar plots showed reductions in pro-inflammatory drivers, notably IL-6 and IL-17 (all p≤0.0384, individual t-tests) while other cytokines remained stable (**Fig. 4D**). In contrast, plasma IL-10 was slightly increased in rewilded mice. Accordingly, regulatory ratios IL-10/IL-6 and IL-10/IL-17 were significantly increased, supporting a systemic anti-inflammatory balance (**Fig. 4E**).

Together, these results indicate that indoor rewilding shifts the pulmonary environment toward enhanced effector readiness while simultaneously promoting systemic immune restraint, offering a dual profile of local vigilance and systemic regulation.

### Alveolar-capillary barrier integrity is preserved under rewilding

We next assessed whether the abovementioned changes translated into measurable disruption of the alveolar-capillary barrier, a critical determinant of gas exchange and respiratory health. Immunoblot analyses of whole lung protein extracts showed comparable abundance of the tight-junction proteins zona occludens 1 (ZO-1), occludin, and claudin-5, as well as of the adherens junction protein vascular-endothelial (VE)-cadherin (**Fig. 5A**). Consistently, albumin concentrations in BALF did not differ significantly across the groups, indicating no detectable protein extravasation into the airspaces (**Fig. 5B**). Thus, despite denser leukocyte traffic and elevated cytokines, the alveolar-capillary barrier remained intact.

**Figure 5.**
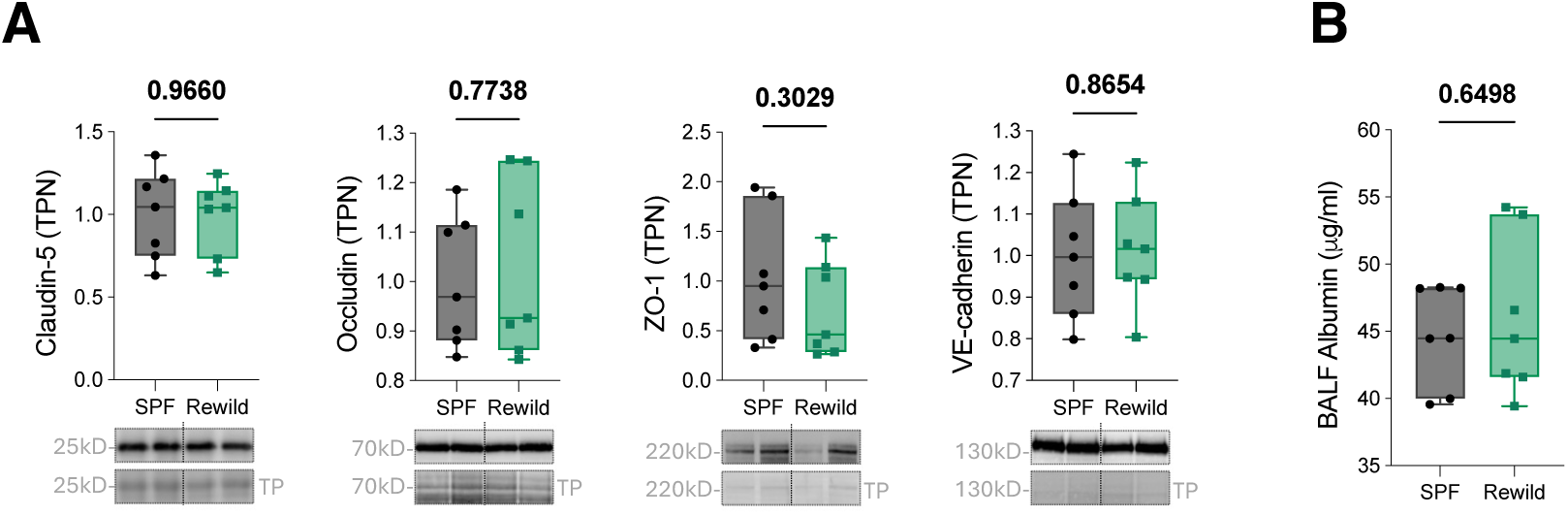
Alveolar-capillary barrier integrity remains preserved under rewilding. **(A)** Immunoblot quantification of junctional proteins claudin-5, occludin, ZO-1, and VE-cadherin from whole-lung homogenates. Representative blots with total protein normalization controls are shown. **(B)** Albumin extravasation into airways assessed by ELISA in BALF, as a proxy for barrier permeability. Means were compared using unpaired two-tailed Student’s t-test. No animals or data points were excluded from analysis.

### Rewilding reshapes the composition and functional potential of the cecal microbiome

Previous studies have shown that mouse rewilding promotes significant changes in microbiota profiles, which could influence, at least in part, immune maturation and responses. Cecal shotgun metagenomics revealed that indoor rewilding markedly restructured bacterial community composition without detectable changes in alpha diversity at all taxa levels (Shannon and observed richness; **Fig. 6A, D**). In contrast, Aitchison (CLR) beta-diversity showed robust separation between SPF and rewilded mice across phylum, genus, and species profiles (PERMANOVA p<0.006), with no evidence that group differences were driven by dispersion (PERMDISP, ns; **Fig. 6B, E, G**). To limit sparsity-driven artifacts, all taxa analysis applied a minimum prevalence threshold (≥4/6 mice per group) prior to differential abundance (DA) testing. DA analysis identified 13 phyla differing between groups (FDR<0.05; **Fig. 6C**) and genus-level shifts consistent with rewilding-driven remodeling, including directional increases in *Akkermansia* and *Bifidobacterium* and decreases in *Lactobacillus* and *Faecalibaculum* (**Fig. 6D**), alongside a subset of differentially abundant genera shown by CLR Z-scores (**Fig. 6F**).

**Figure 6.**
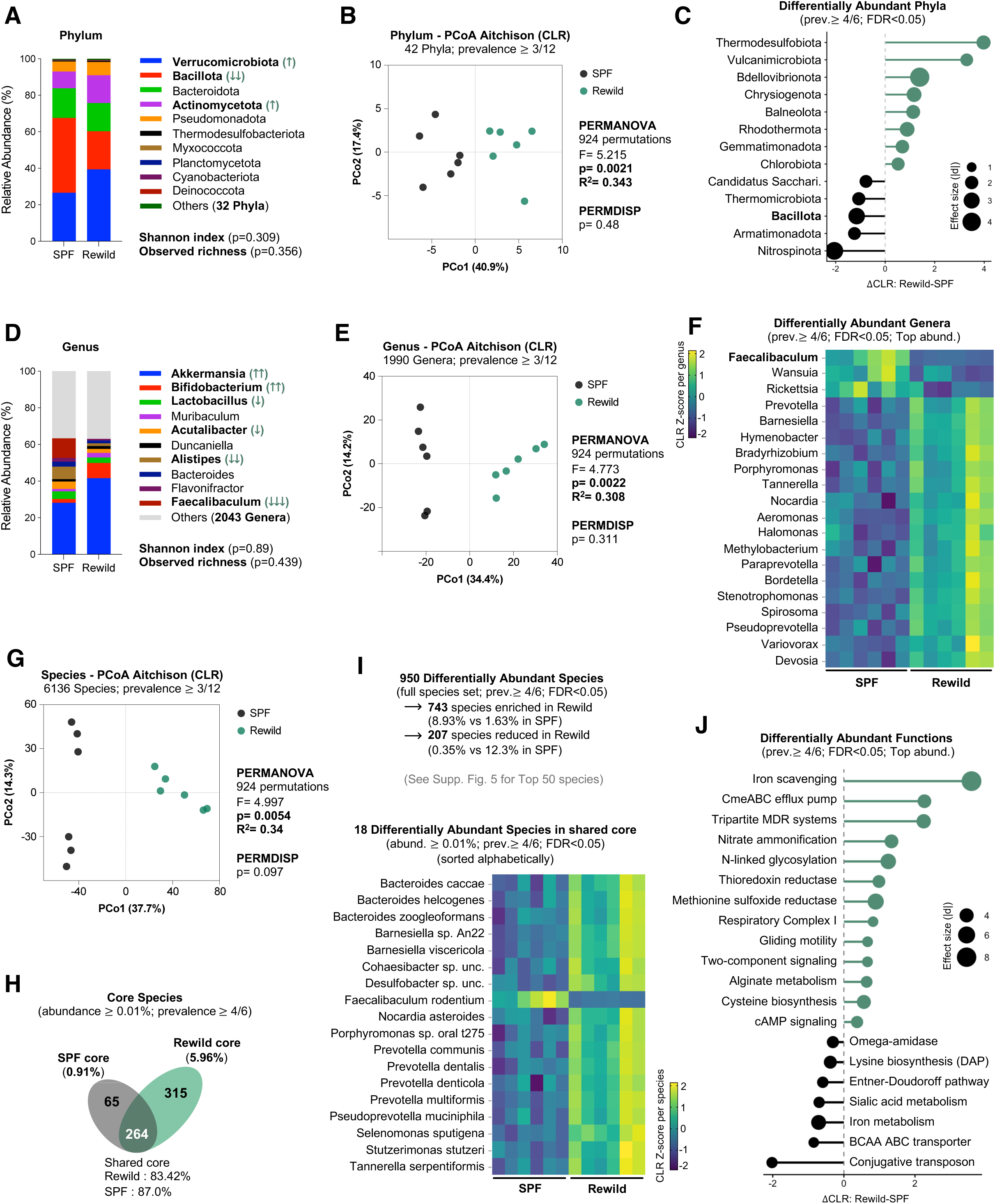
Rewilding reshapes the composition and functional potential of the cecal microbiome. Cecal shotgun metagenomic profiles from SPF and rewilded mice (n=6/group) were analyzed using CLR/Aitchison compositional approaches. **(A, D)** Relative abundance at the phylum and genus levels (top taxa shown; remaining taxa grouped as “Others”); arrows indicate direction and magnitude of mean shifts in Rewild versus SPF (descriptive, not inferential). **(B, E, G)** PCoA of phylum, genus, and species profiles using Aitchison distance on CLR-transformed data (each point represents one mouse; PERMANOVA/PERMDISP results are shown in the panels). **(C, J)** Differentially abundant phyla and SEED subsystem level-3 functions (prevalence filter ≥4/6; FDR<0.05) shown as ΔCLR (Rewild-SPF; 95% CI), where ΔCLR>0 indicates Rewild enrichment and ΔCLR<0 indicates SPF enrichment; point size denotes Cohen’s*-d* effect size. **(F)** Heatmap of differentially abundant genera (prevalence filter ≥4/6; FDR<0.05) with the highest mean relative abundance among significant genera, displayed as per-genus Z-scored CLR across individual mice. **(H)** Venn diagram of core species overlap (core was defined within each group as taxa detected in ≥4/6 and mean relative abundance ≥0.01%); numbers denote species counts and percentages indicate the cumulative relative-abundance contribution of each subset within the corresponding group. **(I)** Species-level DA summary for the full species set and heatmap of the shared-core DA species, shown as per-species Z-scored CLR computed from the full feature set (see **Supp. Fig. 6** for the top 50 species). Unc.: Uncultured.

Core-species analysis showed substantial overlap between housing conditions (264 shared core species) but a larger and more abundant rewild-specific core (315 species; 5.96% cumulative abundance) compared with SPF (65 species; 0.91%; **Fig. 6H**). Across the full species set, 950 taxa were differentially abundant (filter prior to DA: prevalence ≥4/6 and abundance ≥0.01%; FDR<0.05): 743 species enriched in rewilded mice contributed 8.93% cumulative abundance (vs 1.63% in SPF), whereas 207 species reduced in rewilded mice contributed 0.35% (vs 12.3% in SPF; **Fig. 6I**; **Supp. Fig. 6**). Within the shared core, 18 species remained differentially abundant (FDR<0.05; **Fig. 6I**).

These taxonomic shifts were paralleled by functional remodeling, with rewilding enriching pathways related to iron scavenging, efflux/MDR, nitrate ammonification, redox/respiration (e.g., respiratory complex I; thioredoxin/methionine sulfoxide reductases), and signaling/motility, while SPF was relatively enriched for functions including sialic acid metabolism and lysine biosynthesis (**Fig. 6J**). Taken together, these results are consistent with a systemic remodeling of immune maturation associated with environmental enrichment, and delineate candidate ecological and microbial features that could be mechanistically tested in future work.

## DISCUSSION

“Dirty” mouse models have advanced our understanding of how ecological exposures shape the immune system, although their effects beyond immune remodeling, particularly on vital barrier organs physiology remain largely underexplored. The lung, as a continuously exposed barrier organ requiring the integration of mechanical performance, immune vigilance, and tissue integrity, represents a particularly sensitive system to test such effects. In this study, we demonstrate that indoor rewilding restores immune complexity within a relatively short period (90 days) while recalibrating pulmonary biomechanics and systemic homeostasis in SPF-reared adult mice. Accordingly, ecological enrichment extends beyond immunological recalibration to support organ-level functional and tissue resilience.

Our work extends previous advances from “dirty” models such as fecal microbiota transfer (FMT) *via* oral gavage or coprophagy, naturalization protocols, and (outdoor) rewilding approaches that have linked microbial complexity to immune maturation [1,2,15,16,30–33]. In addition to organic materials, our approach incorporates living vegetation and invertebrates within a controlled indoor environment, thereby providing ecological enrichment while maintaining experimental containment. Unlike previous rewilding paradigms, which can expose animals to climate variations, predators, and unpredictable pathogens such as helminths or ectoparasites, the indoor rewilding framework preserves biosecurity and reproducibility and can be readily adapted to existing non-SPF infrastructures. Based on source selection and provenance of organic materials used in this vivarium, the risk of introducing murine-specific pathogens was *a priori* considered minimal. Post-exposure health monitoring detected no external pathogens introduced over the rewilding period (**Supp. Table 4**), supporting the biosafety of our approach. While this post-hoc readout reduces the likelihood of inadvertent pathogen introduction, pre-introduction environmental screening would further strengthen biosafety assurance and reproducibility across sites. Embedding this rewilding paradigm within an accredited animal facility streamlines essential operations such as colony transfer, health surveillance, and longitudinal experimental design, offering a scalable and standardized platform for modeling immune-functional integration studies under semi-natural conditions.

The pulmonary immune remodeling observed in rewilded mice reiterates but also extends prior wildling findings. Airspaces and parenchyma displayed denser leukocyte subsets, with expanded CD4⁺ and CD8⁺ T cells, NKT and NK populations, and elevated immune checkpoint expression (PD-L1/PD-L2) on cDC2s. This configuration supports surveillance and tolerance rather than allergy, as eosinophils and IL-4/IL-5 responses were absent. The absence of eosinophilic infiltration, despite adult-phase exposure, likely reflects the establishment of local regulatory control during the three-month rewilding period, which conditioned tolerance rather than hypersensitivity to environmental cues. The cytokine profile within the lung, marked by TNF, IFN-γ, CXCL1/CXCL2, CCL3/CCL4, and IL-10 upregulation, reflects a vigilant yet balanced immunophenotype. Despite this local activation, systemic cytokine patterns leaned toward anti-inflammatory balance (elevated IL-10/IL-6 and IL-10/IL-17 ratios), consistent with compartmentalized mucosal homeostasis uncommon in SPF mice. These cytokine contrasts were paralleled by increased PECAM-1 and ICAM-1 expression, promoting selective leukocyte trafficking without excessive inflammation [34,35]. This arrangement resembles human mucosal immunophysiology, where immune readiness coexists with structural containment [36–38].

Despite intensified immune activity, alveolar-capillary barrier integrity remained intact. Tight- and adherens-junction proteins (ZO-1, occludin, claudin-5, VE-cadherin) and BALF albumin were unchanged, indicating preserved barrier function and demonstrating that immune trafficking can occur without structural compromise. Notably, in inflammatory models, TNF/ICAM-1 upregulation usually accompanies junctional breakdown [39]. Thus, indoor rewilding promotes a phenotype of regulated tissular resilience which separates immune activation from barrier alteration, likely mediated through coordinated adhesion and checkpoint mechanisms (PECAM-1, PD-L pathways). Clinically, this distinction is critical since barrier dysfunction drives asthma, COPD, and fibrosis.

Functionally, the mechanical consequences of this immune recalibration were mirrored in FlexiVent analysis. Rewilded mice exhibited lower central airway resistance, improved compliance, and blunted methacholine-induced reactivity, hallmarks of more efficient lung mechanics. These enhancements suggest balanced immune-epithelial interactions and tissue maintenance, limiting edema formation and blunting pro-contractile cascades classically linked to AHR. This immune-mechanical homeostasis may explain the paradoxical association of elevated TNF with reduced AHR in rewilded mice. In SPF asthma models, TNF promotes airway hyperresponsiveness, and anti-TNF agents alleviate symptoms [40–43]. Here, concurrent IL-10 elevation and PD-L1/PD-L2 upregulation likely buffer TNF’s contractile signaling, redirecting it toward coordination rather than pathology. This context-specific modulation underscores how immunological immaturity in SPF mice can distort cytokine function and thus therapeutic predictability.

Comparing our findings to previous studies provides further insight. Islam et al. [16] reported that SPF mice colonized with wild-derived gut microbiota showed reduced house-dust-mite (HDM)-induced allergic inflammation and AHR *via* TNF-dampening and metabolic modulation, consistent with our finding of reduced airway reactivity under methacholine. Conversely, Ma et al. [15] found that wild-inherited microbiota amplified Th2 inflammation in HDM-challenged offspring. Unlike that model, our study examined baseline physiology and direct smooth-muscle reactivity, revealing a freer, non-hyperreactive airway state. Together, these contrasts emphasize the role of exposure context (direct *vs* transgenerational, baseline *vs* allergen-challenged) in determining whether rewilding fosters tolerance or pathology.

Furthermore, rewilded mice exhibited systemic adaptations aligned with improved pulmonary physiology, including enhanced neuromuscular performance and reduced plasma EPO despite increased performance, consistent with improved ventilation and tissue oxygenation and reduced hypoxic drive. Stable corticosterone further suggests that indoor rewilding did not induce chronic stress. Together, these changes point to improved energy efficiency and systemic resilience under ecological enrichment.

Unsurprisingly, rewilding also reshaped the cecal microbiome, expanding a rewild-specific core and redistributing species abundance within the shared core, consistent with prior rewilding paradigms [33]. Notably, several rewild-enriched genera (e.g., *Bifidobacterium*, *Barnesiella*) have documented immunomodulatory or colonization-resistance properties, providing a plausible microbial context for downstream host immune changes [44]. Of particular interest is the ecological contribution of diverse organic living materials naturally associated with soil, plants, and water sources, reflected by the presence of canonical environmental taxa such as *Streptomyces* and *Micromonospora* (soil actinobacteria), plant-associated rhizobia including *Bradyrhizobium*/*Mesorhizobium*, and aquatic-associated genera such as *Aeromonas* and *Marinobacter*. Notably, an insect-associated signal was also captured by detecting *Streptomyces formicae*. Whether these contribute to the lung phenotypes observed herein remains to be tested. Interestingly, the species-level bacterial remodeling was accompanied by extensive functional shifts in pathways related to iron acquisition, redox/respiration, transport/efflux, and signaling/motility. Collectively, these coordinated microbial and functional changes provide a valid ecological context for studying immune and physiological recalibration, while preserving barrier integrity and systemic homeostasis.

Traditional laboratory mice exhibit altered physiology that can compromise their translational value in preclinical drug testing. Here, we present a “nature-in-the-lab” vivarium that combines ecological realism with experimental control. By restoring immune-functional integration, this model enhances predictive power for conditions like asthma, COPD, and fibrosis, where immune and mechanical factors intersect, and strengthens preclinical testing for translatable therapies. By exposing laboratory mice to controlled ecological complexity, this system not only enhances animal welfare but also broadens the scope of biomedical research in respiratory physiology, immunology, and beyond. Although it cannot capture all real-world pressures (e.g., seasonal climate variation, predator stress, population dynamics, poly-parasitism), indoor rewilding offers a tractable bridge between SPF housing and more naturalized exposures, complementary to outdoor rewilding. Importantly, while effects were consistent within this design, additional cohorts and replication studies will be important to confirm generalizability.

In conclusion, indoor rewilding establishes a balance between immune vigilance and biomechanical efficiency rarely seen in SPF animals. This environment restores human-like mucosal immune profiles, maintains epithelial and endothelial barrier integrity, and improves respiratory mechanics while reducing airway hyperresponsiveness. Future longitudinal, sex-balanced studies could examine the longevity of these effects, possible sex differences, strain dependence (e.g., BALB/c), and interactions with disease-susceptible genotypes. Whether similar functional findings can be reproduced in other “dirty” models remains to be tested.

## SUPPLEMENTARY MATERIALS

**Supplementary Figure 1. Semi-natural indoor vivarium setup. (A)** Schematic representation of the indoor rewilding vivarium constructed using an adapted, reinforced Coleman™ steel-frame swimming pool (L: 4.88m, W: 3m, H: 1.1m). Enclosures were filled with 30-40 cm of organic soil and enriched with natural materials including hay, growing grass, moss, mixed plants, wood logs, bark, rocks, and chicken manure. Additional live ecological complexity was provided by insects (earthworms, ants, moths, flies, crickets). Animals had continuous access to water stations, plant-cycle lighting (12h/12h), modular wooden shelters, bird seeds, and lab chow presented both in bowls and scattered on the ground to support natural behaviors such as burrowing and foraging. Refreshments of organic material (e.g., hay, manure, bark, moss) occurred every 3-4 weeks to sustain continuous antigenic stimulation. A perimeter safety net and added poles ensured structural stability and containment. Environmental parameters (∼22°C, ∼45%RH, 12:12-hour light/dark cycle) and predator exclusion were maintained under controlled indoor conditions. Day and infrared-night cameras monitored the enclosures and room for potential escape. (designed in BioRender).

**Supplementary Figure 2. Representative gating strategy for BAL lymphocyte subsets.** Sequential gating from Time, FSC/SSC, singlets, live cells, and CD45+ leukocytes, followed by lineage markers (CD90.2, CD4, CD8, CD19, NK1.1, CD49b, CD11b).

**Supplementary Figure 3. Representative gating strategy for lung lymphocyte subsets.** Sequential gating from Time, FSC/SSC, singlets, live cells, and CD45+ leukocytes, followed by lineage markers (CD90.2, CD4, CD8, CD19, NK1.1, CD49b, CD11b).

**Supplementary Figure 4. Representative gating strategy for lung granulocyte subsets.** Sequential gating from Time, FSC/SSC, singlets, live cells, and CD45+ leukocytes, followed by lineage markers (CD11b, CD11c, CCR3, Siglec-F, Ly6G).

**Supplementary Figure 5. Representative gating strategy for lung dendritic cell subsets.** Sequential gating from Time, FSC/SSC, singlets, live cells, and CD45+ leukocytes, followed by conventional DCs markers (CD11b, MHCII, CD11c, CD103, SIRPα, XCR1), with further assessment of PD-L1, PD-L2, and CD200 median fluorescence intensities (MFIs).

**Supplementary Figure 6. Top differentially abundant species identified from the full species set.** (A) Heatmap showing the 50 most abundant differentially abundant species (species-level DA; prevalence ≥4/6; FDR<0.05), displayed as per-species Z-scored CLR values across individual mice (SPF, n=6; Rewild, n=6). Species are listed alphabetically; boldface highlights taxa also detected as differentially abundant within the shared core (Fig. 6I). Unc.: Uncultured.

**Supplementary Table 1.** List of fluorochrome-conjugated antibodies used in flow cytometry panels for lung and BAL leukocyte profiling.

**Supplementary Table 2.** List of forward and reverse primers designed for amplification of murine cytokine, chemokine and housekeeping genes. Primer sequences were used to quantify mRNA expression in Lung tissue by SYBR Green-based qRT-PCR.

**Supplementary Table 3.** List of primary and secondary antibodies used for protein detection by immunoblot. Dilutions used, host species, suppliers, and catalog numbers are indicated.

**Supplementary Table 4.** Pathogen monitoring of SPF and rewilded mice performed by Charles River’s PRIA® panel. The table lists viral, bacterial, and parasitic agents tested in cecal feces; all results were negative, supporting the absence of pathogen introduction during the rewilding period.

**Supplementary Video 1. Semi-natural indoor vivarium setup.** The vivarium was enriched with farmland soil (∼30-40 cm in depth), dried hay, living grass and plants, a small wooden shed, wood chips and bark shedding, rocks and pebbles, mixed chicken and cow manure, fungi, and a variety of invertebrates (worms, ants, grasshoppers, flies), with full exclusion of predators. Pre-introduction pathogen screening of the enrichment substrates was not conducted, because all materials were sourced from human-managed or horticultural environments where the risk of murine-specific pathogens is inherently low. Food and water were provided *ad libitum* at dedicated stations inside the vivarium and monitored daily. In addition to standard lab chow, wild bird seeds were scattered within the enclosure. Refreshments of organic material (e.g., hay, manure, bark, moss) occurred every 3-4 weeks to sustain continuous antigenic stimulation.

**Supplementary Video 2. Behavioral activity of rewilded mice.** Infrared recordings showing spontaneous locomotor and exploratory behaviors of C57BL/6J mice during the three-month indoor rewilding period, illustrating enhanced physical activity and natural behaviors such as food caching and tunnel exploration.

## Notes

### Competing Interest Statement

The authors have declared no competing interest.

### Summary of Updates

The revised manuscript includes cecal-derived shotgun metagenomic profiling, providing species-level and functional insights into how our indoor rewilding paradigm reshapes the gut microbiome. We have also expanded the description of the semi-natural vivarium, and discuss the strengths and limitations of indoor rewilding relative to other dirty-mouse and outdoor paradigms.

