## Supplementary material for "Indoor Rewilding of Laboratory Mice Recalibrates Pulmonary Mucosal Immunity and Mechanics": Revised Supplementary Materials

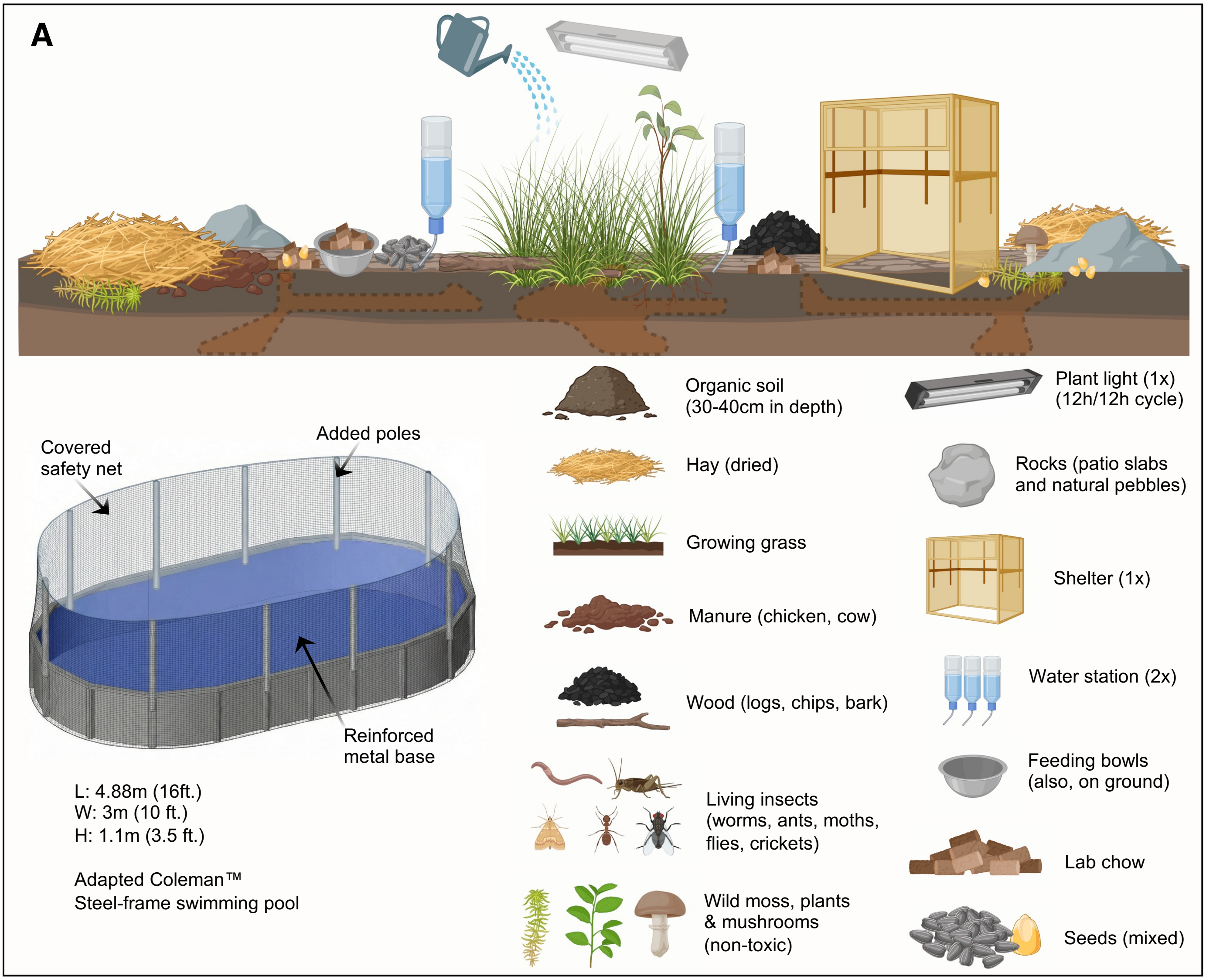


**Supplementary Figure 1. Semi-natural indoor vivarium setup. (A)** Schematic representation of the indoor rewilding vivarium constructed using an adapted, reinforced Coleman™ steel-frame swimming pool (L: 4.88m, W: 3m, H: 1.1m). Enclosures were filled with 30-40 cm of organic soil and enriched with natural materials including hay, growing grass, moss, mixed plants, wood logs, bark, rocks, and chicken manure. Additional live ecological complexity was provided by insects (earthworms, ants, moths, flies, crickets). Animals had continuous access to water stations, plant-cycle lighting (12h/12h), modular wooden shelters, bird seeds, and lab chow presented both in bowls and scattered on the ground to support natural behaviors such as burrowing and foraging. Refreshments of organic material (e.g., hay, manure, bark, moss) occurred every 3-4 weeks to sustain continuous antigenic stimulation. A perimeter safety net and added poles ensured structural stability and containment. Environmental parameters (~22°C, ~45%RH, 12:12-hour light/dark cycle) and predator exclusion were maintained under controlled indoor conditions. Day and infrared-night cameras monitored the enclosures and room for potential escape. (designed in BioRender)

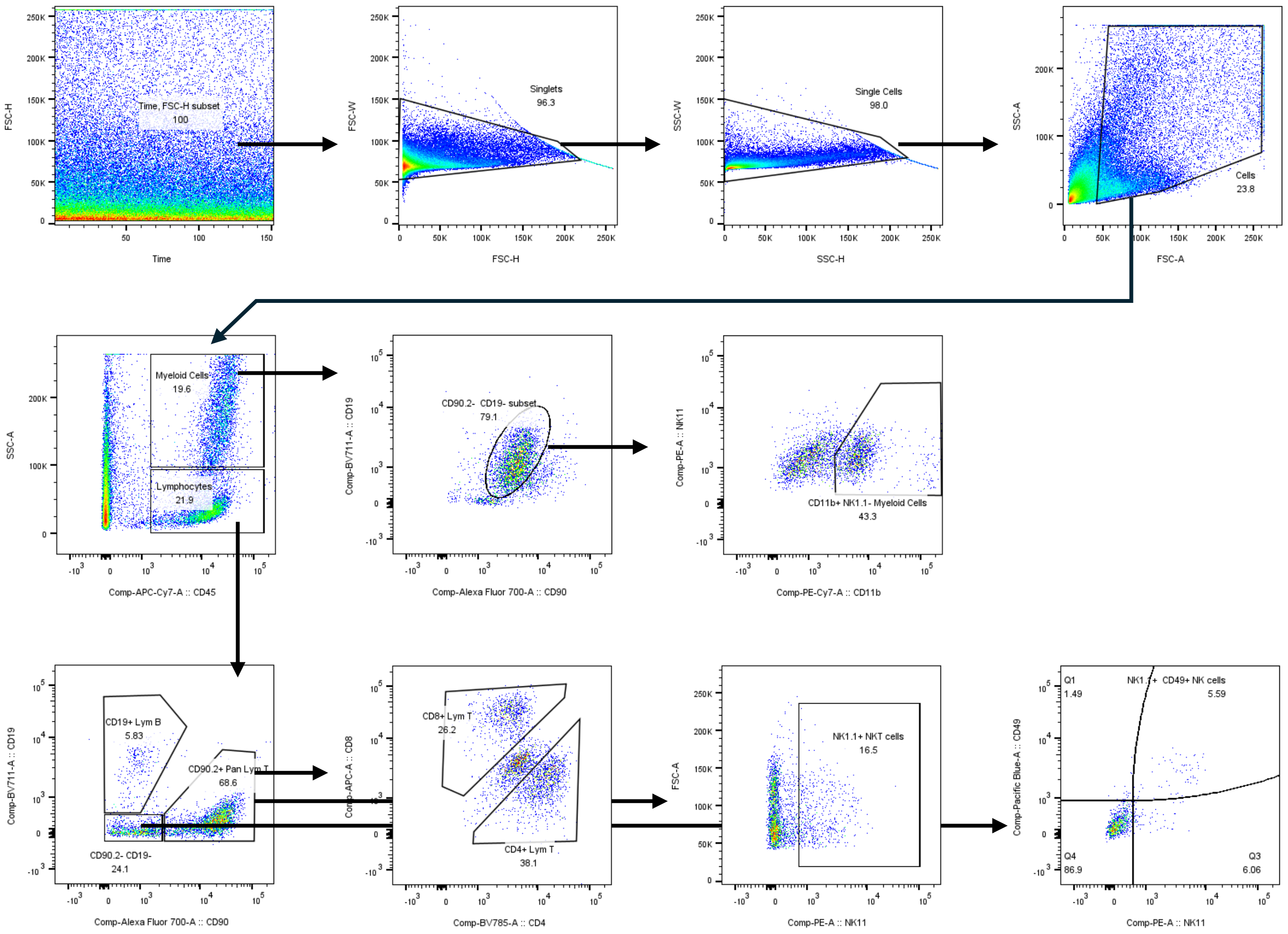


**Supplementary Figure 2. Representative gating strategy for BAL Lymphocyte subsets.** Sequential gating from Time, FSC/SSC, singlets, live cells, and $CD45^{+}$ leukocytes, followed by lineage markers (CD90.2, CD4, CD8, CD19, NK1.1, CD49b, CD11b).

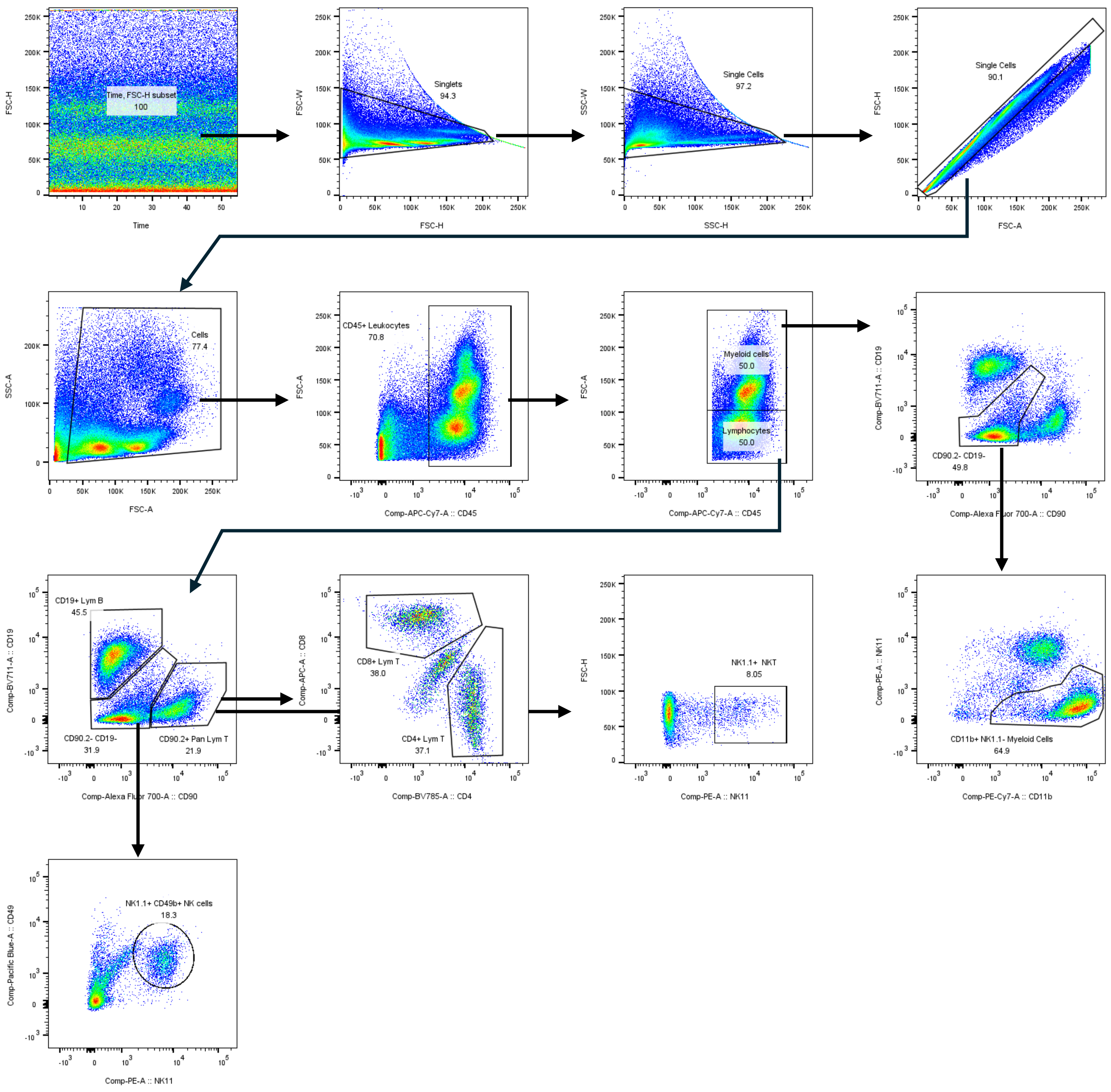


**Supplementary Figure 3. Representative gating strategy for Lung Lymphocyte subsets.** Sequential gating from Time, FSC/SSC, singlets, live cells, and CD45$^{+}$ leukocytes, followed by lineage markers (CD90.2, CD4, CD8, CD19, NK1.1, CD49b, CD11b).

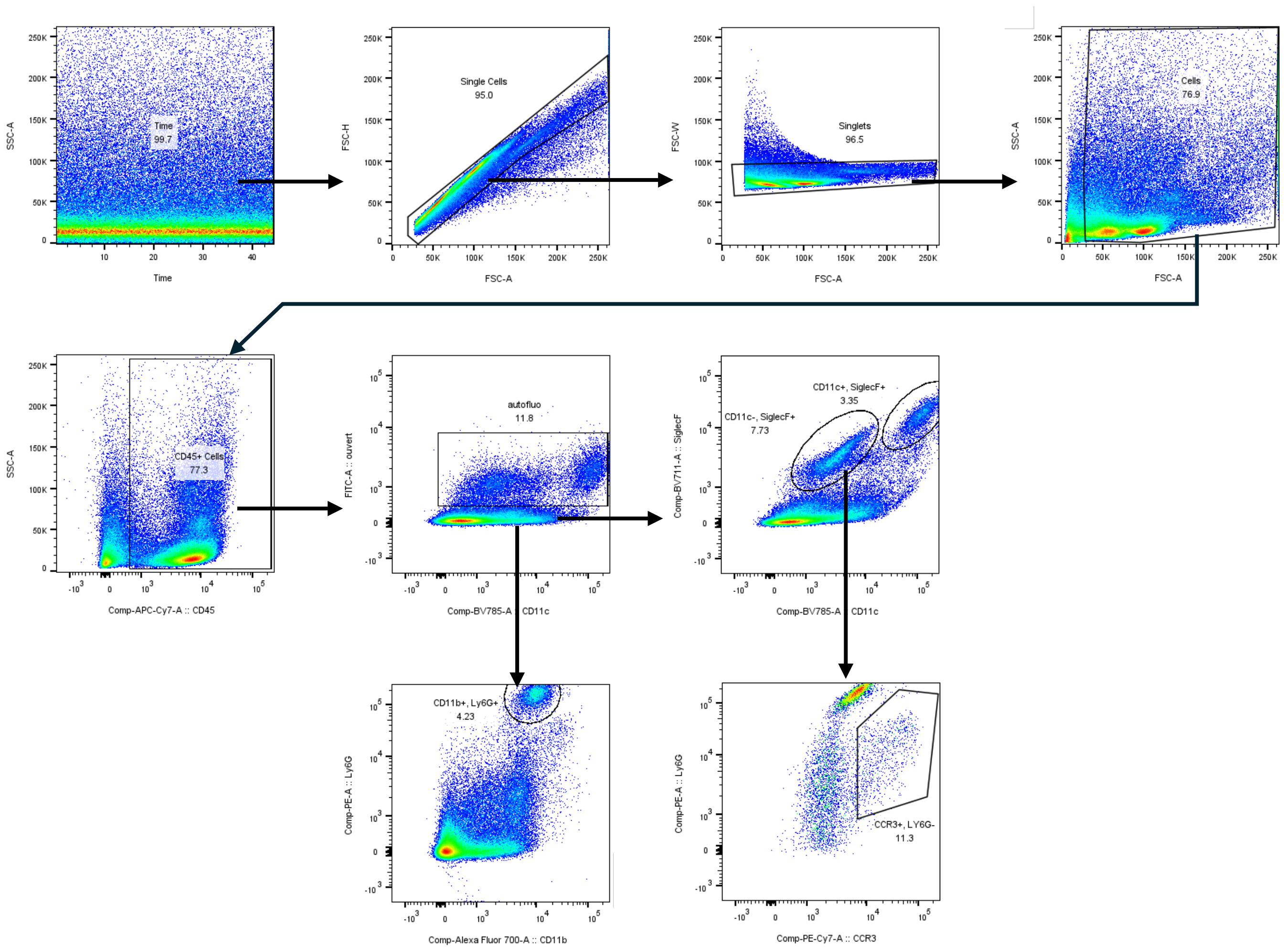


**Supplementary Figure 4. Representative gating strategy for Lung Granulocyte subsets.** Sequential gating from Time, FSC/SSC, singlets, live cells, and $CD45^{+}$ leukocytes, followed by lineage markers (CD11b, CD11c, CCR3, Siglec-F, Ly6G).

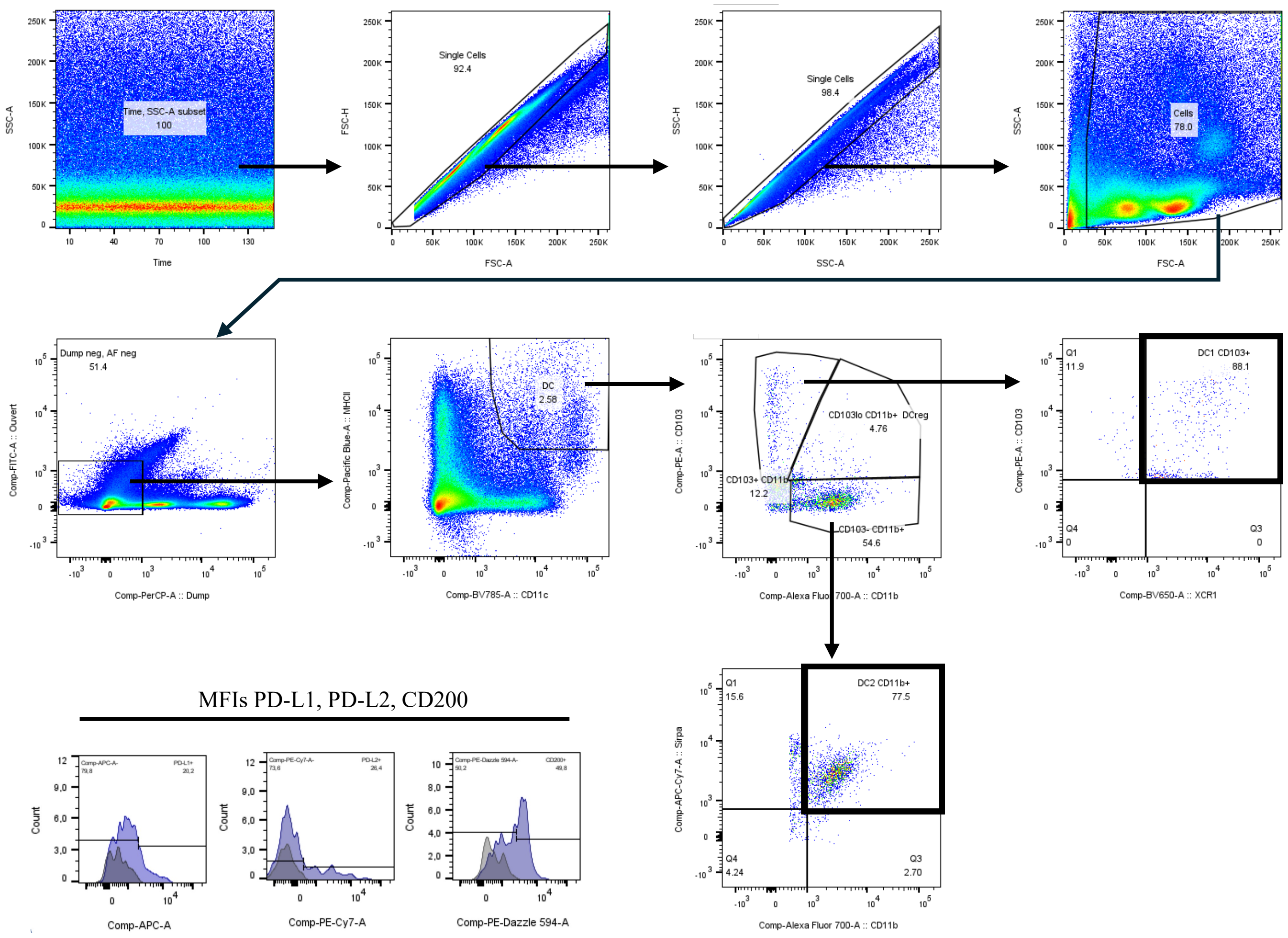


**Supplementary Figure 5. Representative gating strategy for Lung Dendritic Cell subsets.** Sequential gating from Time, FSC/SSC, singlets, live cells, and $CD45^{+}$ leukocytes, followed by conventional DCs markers (CD11b, MHC II, CD11c, CD103, Sirpα, XCR1), with further assessment of PD-L1, PD-L2, and CD200 median fluorescence intensities (MFIs).

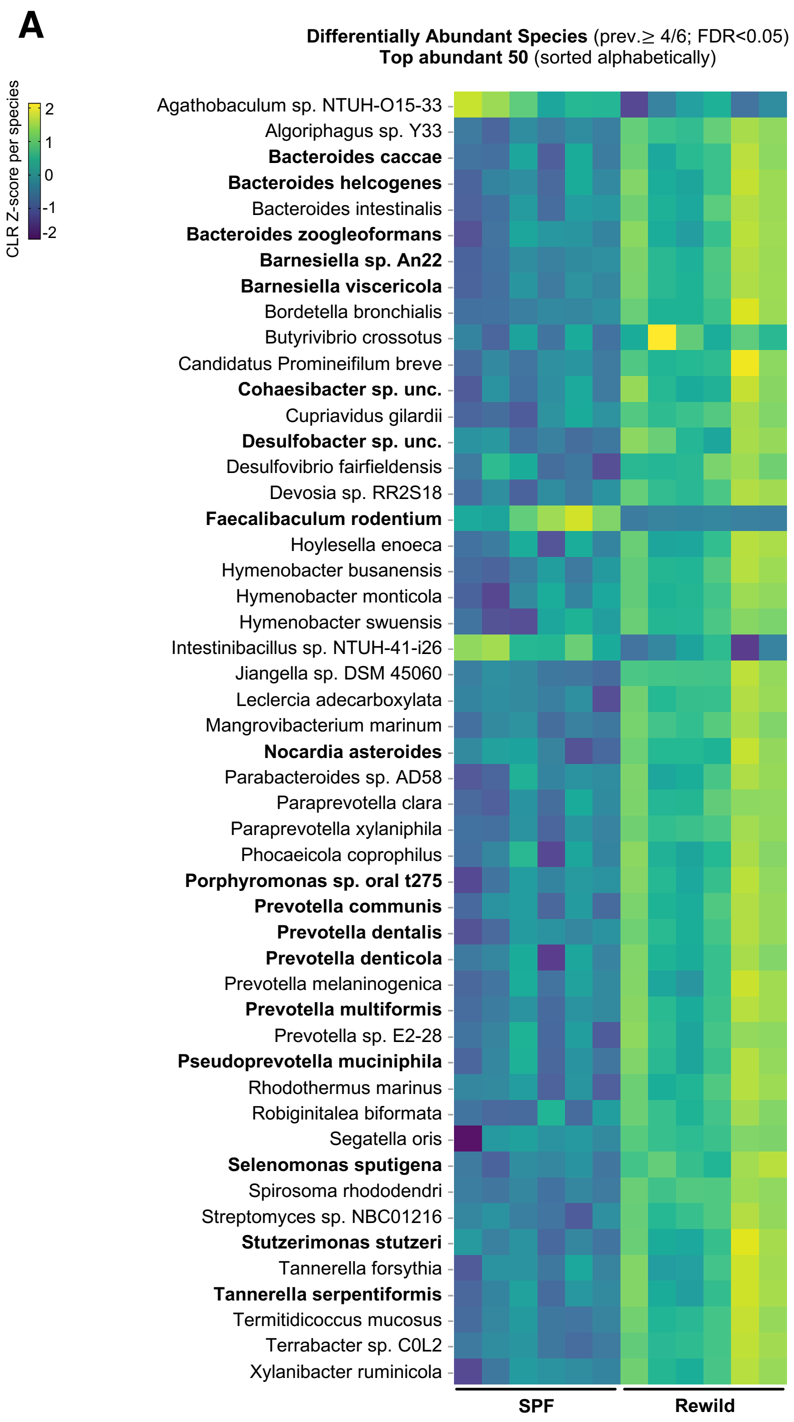


**Supplementary Figure 6. Top differentially abundant species identified from the full species set. (A)** Heatmap showing the 50 most abundant differentially abundant species (species-level DA; prevalence filter ≥4/6; FDR<0.05), displayed as per-species Z-scored CLR values across individual mice (SPF, n=6; Rewild, n=6). Species are listed alphabetically; boldface highlights taxa also detected as differentially abundant within the shared core (Fig. 6I).

**Supplementary Table 1.** List of fluorochrome-conjugated antibodies used in flow cytometry panels for lung and BAL leukocyte profiling.

| Antigen | Fluorochrome | Supplier | # Catalog |
|---|---|---|---|
| Panel for Myeloid Dendritic Cell subsets (Lung) | | | |
| **NK1.1** | Biotin | BioLegend | 108704 |
| **CD19** | Biotin | BioLegend | 115503 |
| **CD90.2** | Biotin | BioLegend | 105304 |
| **MHCII** | PB | BioLegend | 107620 |
| **CD11c** | BV785 | BioLegend | 117335 |
| **SIRPa** | APC-Cy7 | BioLegend | 144018 |
| **CD11b** | AF700 | BioLegend | 101222 |
| **XCR1** | BV650 | BioLegend | 148220 |
| **CD103** | PE | BD Biosciences | 557495 |
| **PD-L1** | APC | BioLegend | 124311 |
| **PD-L2** | PeCy7 | BioLegend | 107214 |
| **CD200** | Pe-Dazzle | BioLegend | 123820 |
| **CD45** | AF700 | BioLegend | 103121 |
| **Streptavidin** | PERCP | BioLegend | 405213 |
| Panel for Lymphocyte subsets (Lung and BAL cells) | | | |
| **CD45** | APC-Cy7 | BioLegend | 103116 |
| **NK1.1** | PE | BioLegend | 108708 |
| **CD49b** | PB | BioLegend | 108917 |
| **CD11b** | PE-Cy7 | BioLegend | 101215 |
| **CD90.2** | AF700 | BioLegend | 105320 |
| **CD8** | APC | BioLegend | 100711 |
| **CD4** | BV785 | BioLegend | 100551 |
| **CD19** | BV711 | BioLegend | 115555 |
| **FCBlock** | - | BioLegend | 103116 |
| Panel for Granulocyte subsets (Lung) | | | |
| **CD45** | APC-Cy7 | BioLegend | 103116 |
| **CD11b** | AF700 | BioLegend | 101222 |
| **CCR3** | PeCy7 | BioLegend | 144514 |
| **Siglec F** | BV711 | BioLegend | 740764 |
| **CD11c** | BV785 | BioLegend | 117335 |
| **Ly6G** | PE | BioLegend | 127607 |

**Supplementary Table 2.** List of forward and reverse primers designed for amplification of murine cytokine, chemokine and housekeeping genes. Primer sequences were used to quantify mRNA expression in Lung tissue by SYBR Green-based qRT-PCR.

| | **Forward** | **Reverse** |
|---|---|---|
| Target murine genes | | |
| **TNF-α** | CAGACCCTCACACTCAGATCAT | TTGCTACGACGTGGGCTAC |
| **CCL4** | TTTCTCTTACACCTCCCGGC | TCTTTTGGTCAGGAATACCACAG |
| **CXCL10** | GCCCACGTGTTGAGATCATT | TTAAGGAGCCCTTTTAGACCTTTT |
| **IL-10** | GCTGTCATCGATTTCTCCCCT | AGACACCTTGGTCTTGGAGCTTA |
| **IL-4** | TCAACCCCCAGCTAGTTGTC | ACTCTCTGTGGTGTTCTTCGTT |
| **IL-5** | TTGACCGCCAAAAAGAGAAGTG | TGCCCACTCTGTACTCATCA |
| **IL-6** | GGAGCCCACCAAGAACGATA | CAACTGGATGGAAGTCTCTTGC |
| **IFN-γ** | TGGCTGTTTCTGGCTGTTAC | GGATTTTCATGTCACCATCCTTTTG |
| **CXCL15** | CCATGGGTGAAGGCTACTGT | GTCCTCAGGTAGGAACCTGTTAG |
| **TGF-β1** | CGTGGAAATCAACGGGATCA | TAGTTGGTATCCAGGGCTCTC |
| Housekeeping murine genes | | |
| **RPLP0** | TCCTCGTTGGAGTGACATCG | CTGTCTTCCCTGGGCATCAC |
| **RPLP2** | TCGCACGCGTGAGCAT | CTGATGACCTTGTTGAGCCG |

**Supplementary Table 3.** List of primary and secondary antibodies used for protein detection by immunoblot. Dilutions used, host species, suppliers, and catalog numbers are indicated.

| Protein | Dilution | Species | Supplier | # Catalog |
|---|---|---|---|---|
| Primary antibodies | | | | |
| **PECAM-1** | 1/1000 | Mouse | Thermo Fisher | MA5-13188 |
| **ICAM-1** | 1/250 | Mouse | Thermo Fisher | MA5407 |
| **VCAM-1** | 1/1000 | Rabbit | Abcam | AB134047 |
| **ZO-1** | 1/1000 | Rabbit | Thermo Fisher | 61-7300 |
| **Claudin-5** | 1/1000 | Mouse | Thermo Fisher | 35-2500 |
| **Occludin** | 1/500 | Rabbit | Thermo Fisher | 71-1500 |
| **VE-cadherin** | 1/1000 | Rabbit | Abcam | AB33168 |
| Secondary antibodies | | | | |
| **HRP-conjugated IgG (HC+LC)** | 1/5000 | Goat anti-mouse | Jackson laboratories | 115-035-003 |
| **HRP-conjugated IgG (HC+LC)** | 1/5000 | Goat anti-rabbit | Jackson laboratories | 111-035-144 |

**Supplementary Table 4.** Pathogen monitoring of SPF and rewilded mice performed by Charles River's PRIA® panel. The table lists viral, bacterial, and parasitic agents tested in cecal contents; all results were negative, confirming the absence of pathogen introduction during the rewilding period.

| Agents | SPF mice | Rewild mice |
|---|---|---|
| Viruses | | |
| **LCMV** | Negative | Negative |
| **MAV 1 & 2** | Negative | Negative |
| **MHV** | Negative | Negative |
| **MNV** | Negative | Negative |
| **Mousepox (Ectromelia)** | Negative | Negative |
| **Mouse Parvovirus (MPV/MVM)** | Negative | Negative |
| **MRV (EDIM)** | Negative | Negative |
| **PVM** | Negative | Negative |
| **REO** | Negative | Negative |
| **SEND** | Negative | Negative |
| **TMEV/GDVII** | Negative | Negative |
| **Hantavirus** | Negative | Negative |
| Pathogenic Bacteria | | |
| **Beta Strep Grp A** | Negative | Negative |
| **Beta Strep Grp B** | Negative | Negative |
| **Beta Strep Grp C** | Negative | Negative |
| **Beta Strep Grp G** | Negative | Negative |
| **B. bronchiseptica** | Negative | Negative |
| **Campylobacter Genus** | Negative | Negative |
| **C. kutscheri** | Negative | Negative |
| **C. piliforme** | Negative | Negative |
| **Ps. aeruginosa** | Negative | Negative |
| **Salmonella Genus** | Negative | Negative |
| **S. aureus** | Negative | Negative |
| **S. moniliformis** | Negative | Negative |
| **S. pneumoniae** | Negative | Negative |
| Commensal / Opportunistic Bacteria | | |
| **B. pseudohinzii** | Negative | Negative |
| **C. bovis** | Negative | Negative |
| **Filobacterium rodentium (CAR Bacillus)** | Negative | Negative |
| **C. rodentium** | Negative | Negative |
| **K. oxytoca** | Negative | Negative |
| **K. pneumoniae** | Negative | Negative |
| **Helicobacter genus** | Negative | Negative |
| **M. pulmonis** | Negative | Negative |

| | | |
|---|---|---|
| **R. heylii** | Negative | Negative |
| **R. pneumotropicus** | Negative | Negative |
| **P. mirabilis** | Negative | Negative |
| Parasites / Protozoa / Helminths | | |
| **Cryptosporidium** | Negative | Negative |
| **Demodex** | Negative | Negative |
| **Entamoeba** | Negative | Negative |
| **Giardia** | Negative | Negative |
| **Mite** | Negative | Negative |
| **Pinworm** | Negative | Negative |
| **Pneumocystis** | Negative | Negative |
| **Spironucleus muris** | Negative | Negative |
| **Tritrichomonas genus** | Negative | Negative |
